# ZBTB7A regulates MDD-specific chromatin signatures and astrocyte-mediated stress vulnerability in orbitofrontal cortex

**DOI:** 10.1101/2023.05.04.539425

**Authors:** Sasha L. Fulton, Jaroslav Bendl, Isabel Gameiro-Ros, John F. Fullard, Amni Al-Kachak, Ashley E. Lepack, Andrew F. Stewart, Sumnima Singh, Wolfram C. Poller, Ryan M. Bastle, Mads E. Hauberg, Amanda K. Fakira, Min Chen, Romain Durand-de Cuttoli, Flurin Cathomas, Aarthi Ramakrishnan, Kelly Gleason, Li Shen, Carol A. Tamminga, Ana Milosevic, Scott J. Russo, Filip Swirski, Robert D. Blitzer, Paul A. Slesinger, Panos Roussos, Ian Maze

## Abstract

Hyperexcitability in the orbitofrontal cortex (OFC) is a key clinical feature of anhedonic domains of Major Depressive Disorder (MDD). However, the cellular and molecular substrates underlying this dysfunction remain unknown. Here, cell-population-specific chromatin accessibility profiling in human OFC unexpectedly mapped genetic risk for MDD exclusively to non-neuronal cells, and transcriptomic analyses revealed significant glial dysregulation in this region. Characterization of MDD-specific cis-regulatory elements identified ZBTB7A – a transcriptional regulator of astrocyte reactivity – as an important mediator of MDD-specific chromatin accessibility and gene expression. Genetic manipulations in mouse OFC demonstrated that astrocytic Zbtb7a is both necessary and sufficient to promote behavioral deficits, cell-type-specific transcriptional and chromatin profiles, and OFC neuronal hyperexcitability induced by chronic stress – a major risk factor for MDD. These data thus highlight a critical role for OFC astrocytes in stress vulnerability and pinpoint ZBTB7A as a key dysregulated factor in MDD that mediates maladaptive astrocytic functions driving OFC hyperexcitability.

## Introduction

Major Depressive Disorder (MDD) is a leading cause of disability worldwide^1^ and involves corticolimbic network disruptions associated with recurrent episodes of negative affect, cognitive impairment, somatic deficits, and anhedonia^2–5^. Although relatively understudied in human MDD, the orbitofrontal cortex (OFC) processes affective valence and motivational value in humans, monkeys, and rodents – making it a key prefrontal area involved in the anhedonic symptomatic domains of MDD (i.e., loss of pleasure or motivation)^2^. Functional imaging studies have consistently identified significant OFC changes in MDD patients and demonstrated that OFC hyperactivity correlates with the severity of anhedonic and negative rumination symptoms, suicidality, antidepressant treatment responses, and pathogenic trajectories of the disorder^6–16^. A recent RNA sequencing study that profiled multiple brain regions in human MDD identified the OFC as a region displaying the highest number of differentially expressed genes in female patients, and second highest overall^17^, with distinct alterations in gene expression programs identified in comparison to other prefrontal cortical areas. Despite the OFC’s clear involvement in MDD, the molecular and cellular substrates underlying these functional alterations remain poorly understood.

Disease-related cellular phenotypes are determined by spatiotemporally precise gene expression programs induced by transcription factors (TFs) that interact with their corresponding cis-regulatory DNA elements (CREs) in a cell-type-specific manner^18–21^. Chromatin accessibility profiling can be used to identify the full repertoire of active CREs within a given cell-type, and is thus an essential step towards understanding the regulatory drivers of disease pathology. Here, using FANS-coupled ATAC-seq in human postmortem OFC, we found that both genetic risk variants for MDD and MDD-specific CREs were localized to non-neuronal cell populations (primarily glia). We further found that MDD-specific CREs were significantly enriched for binding sites of the chromatin remodeler ZBTB7A, a putative regulator of astrocyte reactivity that was upregulated in MDD OFC and was found to regulate the expression of MDD-specific CRE target genes.

Extending these studies to mice, we found that Zbtb7a is upregulated following chronic stress exposures (a major risk factor for MDD in humans) in astrocytes specifically, and not in microglia or neurons. We further found that Zbtb7a activity in OFC astrocytes is both necessary and sufficient for behavioral stress responsivity using bidirectional, astrocyte-specific Zbtb7a manipulations in preclinical mouse models of stress. Cell-type-specific ATAC-seq and RNA-seq revealed that Zbtb7a mediates chromatin accessibility in astrocytes to promote aberrant gene expression programs related to astrocyte reactivity, including increased inflammatory signaling and impaired synaptic regulation, which led to cell non-autonomous disruption of glutamate signaling pathways in OFC neurons. Furthermore, using electrophysiological recordings and chemogenetic manipulations, we found that Zbtb7a-mediated astrocyte reactivity promotes OFC neuronal hyperexcitability in response to a mild subthreshold stressor, and that this increased OFC excitability mediates maladaptive social avoidance behaviors following chronic stress. In sum, the results of this cross-species study link stress-induced increases in Zbtb7a expression, similar to that observed in human MDD, with astrocyte reactivity and OFC neuronal hyperexcitability, revealing an important mechanism of stress-induced behavioral deficits related to MDD.

## Results

### Chromatin accessibility profiling in neuronal vs. non-neuronal cells of OFC identifies glial-specific regulatory signatures of human MDD

To first investigate gene expression alterations that are associated with MDD diagnosis in human OFC, we performed bulk RNA-seq on postmortem OFC tissues from 20 MDD cases vs. 19 matched healthy controls (**Figure 1A, Figure S1A**). While neuronal hyperactivity is a well-characterized clinical feature of MDD OFC pathology, both differential expression analysis and weighted gene correlation network analysis (WGCNA)^22^ implicated robust alterations in glial cell function and inflammatory responses in MDD, suggesting a key role for non-neuronal cell dysregulation in this region (**Figure 1B-C, Figure S1B-D**). To assess distinct patterns of chromatin accessibility in neuronal vs. non-neuronal (primarily glial) nuclei of human MDD OFC, we implemented FANS (*F*luorescence-*A*ctivated *N*uclear *S*orting) coupled with ATAC-seq (*A*ssay for *T*ransposase-*A*ccessible *C*hromatin followed by Sequencing) on nuclear preparations obtained from these 20 MDD cases vs. 19 matched healthy controls (**Table S1**). We performed extensive quality control assessments of the ATAC-seq libraries to yield a total of 70 high quality sample libraries (**Figure S1E-N**, and **Table S2**). To define the regulatory programs that specify each cell population, we identified active Open Chromatin Regions (OCRs) in neuronal and non-neuronal samples, which accounted for 4.79% and 2.65% of the genome, respectively (**Figure 1D-E**). Using a curated reference dataset^23^, we confirmed that OCR sets in the FAN-sorted populations displayed expected cell-type enrichment patterns (**Figure 1F**). In accordance with previous findings^20^, neuronal OCRs were found to be more distal to transcription start sites (TSSs) compared to non-neuronal OCRs, reflecting a more complex regulatory scheme and higher levels of functional diversity among neuronal subtypes (**Figure 1F**, **Figure S1F**). Because the majority of genetic variants that influence human disease are located within non-coding regulatory regions of the genome^20^, we next investigated the enrichment of common risk variants for MDD in the detected OCR datasets. We calculated the heritability coefficient^24^ for each set of OCRs, stratified by genomic context (**Figure 1G**), and identified significant enrichment for MDD-associated genetic variants only in non-neuronal-specific promoter OCRs, but not in any of the neuronal OCR sets^25^ (**Figure 1G**). These findings indicate that active regulatory elements within non-neuronal OFC cells are relevant to the genetic risk for affective disorders.

**Fig. 1.**
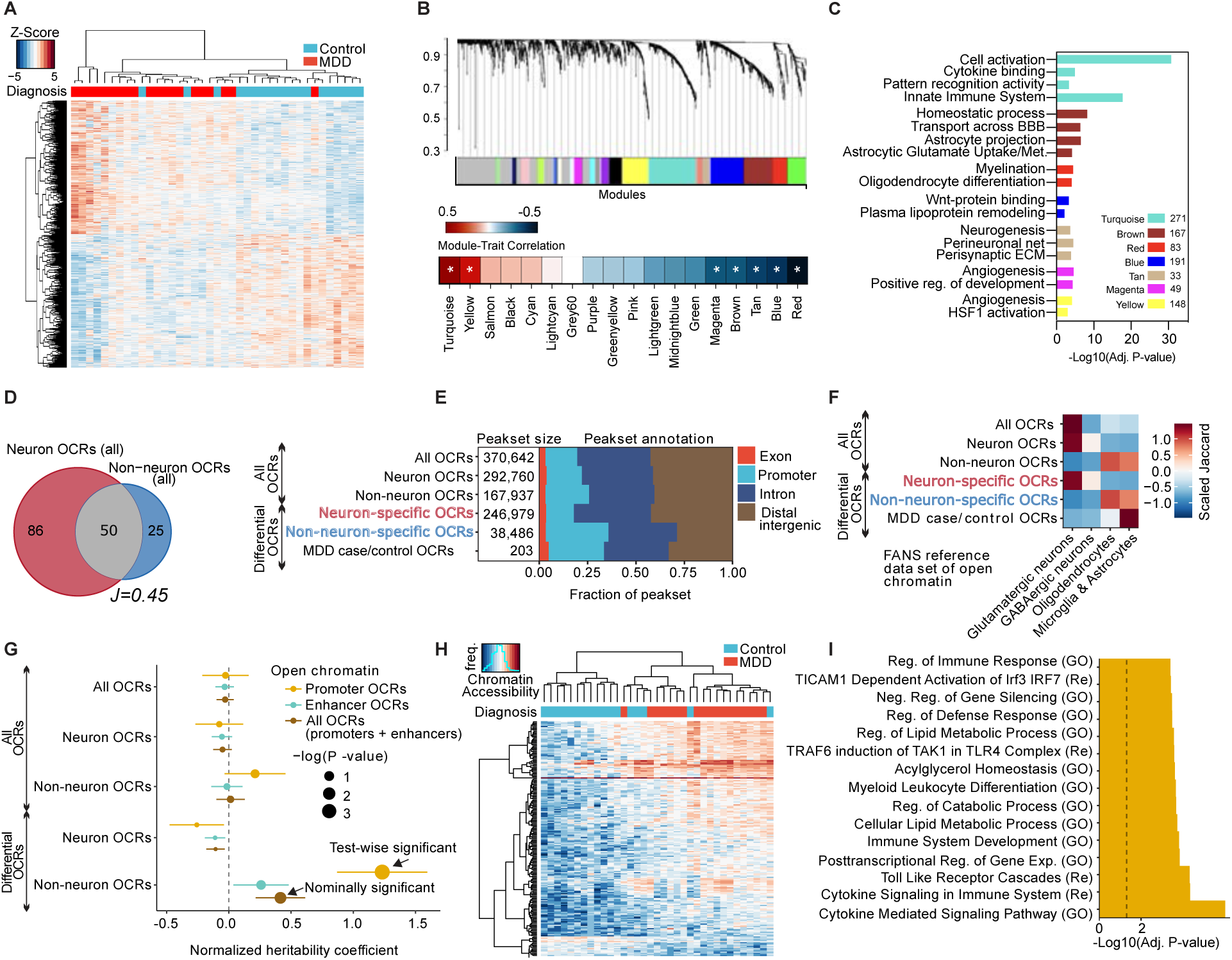
Chromatin accessibility profiling in neuronal vs. non-neuronal cells identifies glial regulatory signatures of human MDD in OFC. (**A**) Clustering of MDD case and control samples at 1412 differentially expressed (DE) genes (rows, FDR < 0.1). (**B**) The co-expression modules identified by weighted gene correlation network analysis (WGCNA) [top] and heatmap of co-expression module correlation with MDD trait. * indicates Adj. P <. 05 significance of correlation. (**C**) Gene Ontology (GO) analysis for genes in significant co-expression modules. (**D**) Venn diagram of shared and distinct open chromatin between neuronal and non-neuronal samples. Numbers indicate megabases of OCRs, “*J*” indicates the Jaccard index. (**E**) Proportions of all and differential OCRs stratified by genomic context. (**F**) Overlap of all and differential OCRs with a reference study of lineage-specific brain open chromatin atlas^61^. (**G**) Enrichment of common genetic variants in MDD^23^ with all and differential OCRs when assayed by LD-score regression. Sets of OCRs were further stratified by genomic context to “Promoter OCRs” overlapping the 3kb window around TSS and “Enhancer OCRs”. (**H**) Clustering of MDD case and control non-neuronal samples at 203 differentially accessible OCRs (rows). (**I**) Overlap between gene sets representing biological processes and pathways with the set of 203 differentially accessible OCRs between MDD cases and control. Top 15 enriched pathways are shown (BH-adjusted p-value < 0.05). Dashed line indicates nominal significance. “GO”: gene ontology, “Re”: REACTOME.

We next assessed differential accessibility at OCRs in each nuclei population to identify putative CREs that were specific to MDD-diagnosis. Consistent with both our RNA-seq analysis and heritability coefficient calculations, we observed differential chromatin accessibility between MDD vs. controls only in non-neuronal OCRs (203 CREs, **Figure 1H**, **Figure S1N-P, Table S3**). We also observed significant correlations between MDD-OCRs and bulk RNA-seq signatures from this same patient cohort (**Figure S1Q)**. Inflammatory gene targets associated with these MDD-specific CREs were found to display significant expression changes in FAN-sorted non-neuronal nuclei from MDD vs. control subjects, such as lower levels of Nuclear Corepressor 2 (*NCOR2),* which was recently identified as a negative regulator of astrocyte-specific reactivity pathways^26^ (**Figure S1R)**. Finally, to characterize the biological processes regulated by MDD-specific CREs in glia, we performed gene set enrichment analysis (GSEA)^27^, which revealed significant changes in pathways associated with glial activation, including NF-kB-induced inflammation, cytokine-mediated cascades, lipid metabolism, and toll-like receptor signaling^28–30^ (**Figure 1I**, **Table S4).** Together, these data demonstrate that MDD-specific CREs mediate cellular stress responses that are known to be disrupted in MDD^30–32^, and converge with previous reported evidence that glial inflammatory stress pathways play a role in the pathophysiology of MDD, particularly in OFC^33–36^.

### Identification of a key transcription factor regulating MDD-specific OCRs: ZBTB7A

To identify potential transcriptional regulators of these MDD-specific CREs, we implemented TF motif discovery analysis^37^, and identified a motif that was significantly enriched (57 motif occurrences out of 202 MDD-specific CREs; **Figure 2A**). In order to characterize the functional role of this enriched regulatory motif, we performed gene ontology (GO) analysis^38^, which revealed significant associations between this enriched motif and gene targets involved in the regulation of inflammatory response (e.g. cytokine pathways and NF-kB cascades) (**Figure 2B**), confirming that this motif is involved in the same regulatory processes that are enriched in MDD-specific OCRs (**Figure 1I**). Out of the top five candidate TFs with binding motifs that matched the enriched motif sequence, only one of these candidates was expressed at detectable levels in human brain and was also dysregulated between MDD and controls: ZBTB7A (**Figure 2C-2D**), which displayed significant upregulation in MDD OFC at both the mRNA and protein levels (**Figure 2E-F** and **Figure S2B**). These findings that are in accordance with previous profiling studies^39^. ZBTB7A is a chromatin regulatory factor with pleiotropic effects (both repressive and activating) and has been shown to coordinate alterations in chromatin structure that are necessary for NF-kB dependent inflammatory gene expression in the context of several types of cancers (notably gliomas) and inflammatory conditions^40^. However, its contributions to psychiatric disease have not yet been explored^41^.

**Fig. 2.**
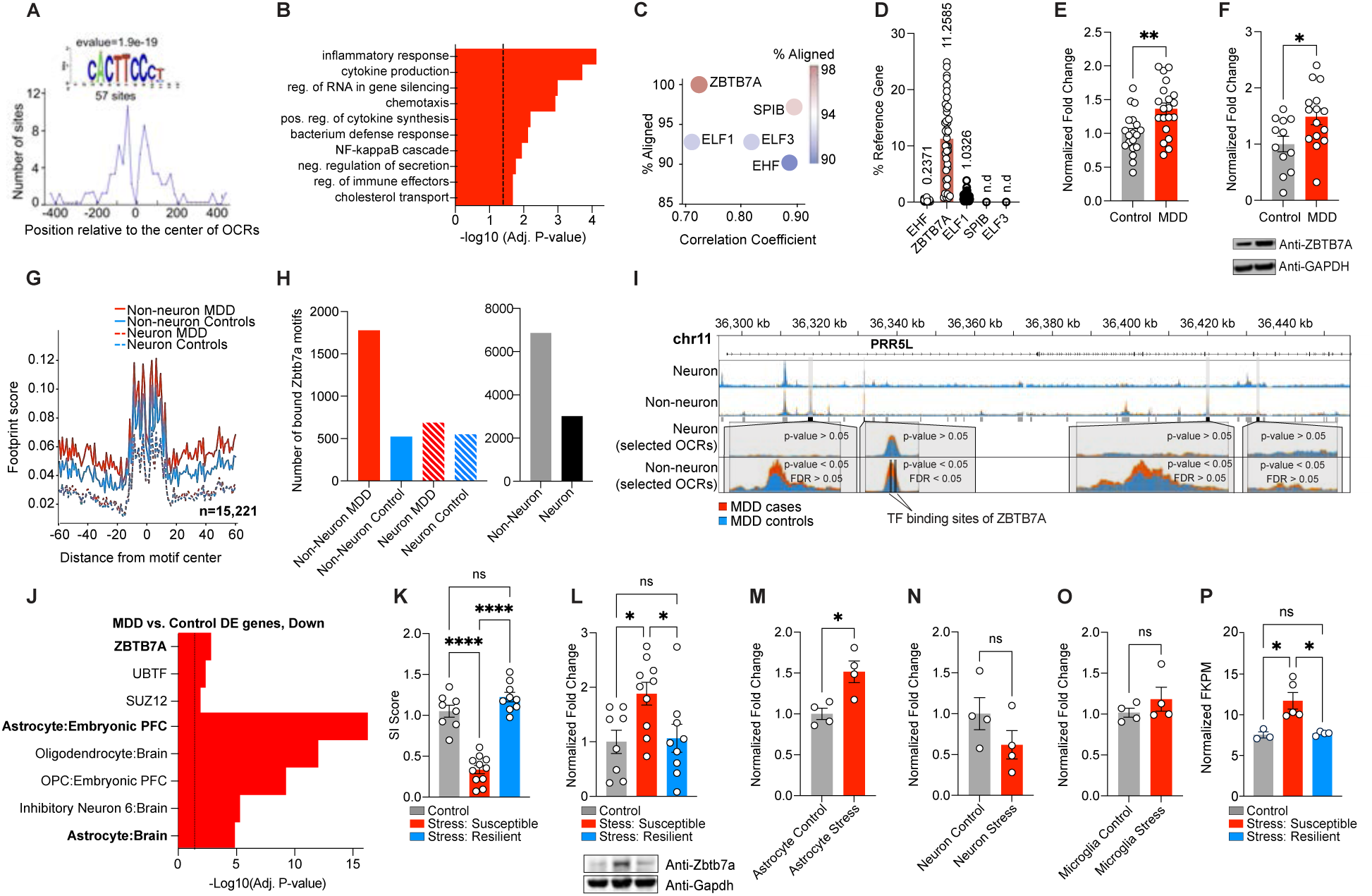
Identification of ZBTB7A as a key transcription factor regulating MDD-specific OCRs. (**A**) Distribution of the discovered motif that is significantly enriched (e-value = 1.9e-19) in MDD-specific OCRs. (**B**) GO BP terms from MEME-GoMo, based on gene targets of regulatory regions containing the discovered motif. Top 10 most significant terms are shown (BH-adjusted p-value < 0.05). Dashed line indicates p = 0.05 significance. (**C**) Correlation coefficients for TF candidate recognition motifs against discovered motif (x-axis), and percent alignment between TF candidate recognition motifs with discovered motif (y-axis and color key) (**D**) Percent expression of TF candidate genes (CT value) over reference gene (*HPRT1*). “n.d.” indicates not detected (**E**) Normalized fold change of *ZBTB7A* transcripts in bulk OFC postmortem human tissues from MDD (n = 20) vs. control (n = 19) samples. Student’s two-tailed t-test [t_37_ = 3.215, **p = 0.0027] (**F**) Normalized fold change of *ZBTB7A* protein in bulk OFC postmortem human tissues from MDD (n = 15) vs. control (n = 12) samples. Student’s two-tailed t-test [t_25_ = 2.441, *p = 0.0221] (**G**) Aggregated footprint scores across ZBTB7A transcription factor binding sites that are bound in either MDD or control samples of neuronal or non-neuronal cells. Note that the effect of Tn5 transposase bias is not fully corrected, resulting into unsmoothed signal. (**H**) Bar graphs for number of bound ZBTB7A TFBS detected exclusively in MDD case or control samples from neuronal and non-neuronal cells (left) and exclusively in non-neuronal and neuronal populations (mixed MDD/control (right). (**I**) Representative pile-up traces of cell specific ATAC-seq signal overlapping PRR5L gene. Four OCRs, all being dysregulated between MDD cases and controls (p-value < 0.05) in non-neuronal cells, are highlighted. The most significantly dysregulated OCR (FDR<0.05) overlaps two transcription factor binding sites of ZBTB7A. (**J**) GO analysis with CellMarker Augmented Database^25^ and CHEA ENCODE Consensus database^108^ for genes in the set of downregulated DE genes from human MDD RNA-seq. (**K**) Social interaction ratio score for control (n = 8) vs. chronic stress: susceptible (n = 11) vs. chronic stress: resilient mouse (n = 9) groups. 1-way ANOVA [F_2,25_ = 66.99], followed by Tukey’s MC test: control vs. stress susceptible ****p=<.0001, stress susceptible vs. stress resilient ****p=<.0001, control vs. stress resilient ns, p = .151. (**L**) Normalized fold change protein expression of Zbtb7a in mouse OFC bulk tissues collected from control vs. chronic stress: susceptible vs. chronic stress: resilient mouse groups. 1-way ANOVA [F_2,24_ = 4.883], followed by Tukey’s MC test: control vs. stress susceptible *p = 0.03, stress susceptible vs. stress resilient *p = 0.039, control vs. stress resilient ns, p = 0.979. (**M**) Normalized fold change *Zbtb7a* mRNA expression in MACs-isolated astrocytes from chronically stressed OFC mouse tissues vs. control (n = 4/group). Two-tailed Student’s t-test [t_6_ = 3.458]. *p = 0.013. (**N**) Normalized fold change *Zbtb7a* mRNA expression in MACs-isolated neurons from chronically stressed OFC mouse tissues vs. control (n = 4/group). Two-tailed Student’s t-test [t_6_ = 1.454]. ns, p = 0.196. (**O**) Normalized fold change *Zbtb7a* mRNA expression in negative cell fraction post MACs-isolation of astrocytes and neurons, which is enriched for microglia, from chronically stressed OFC mouse tissues vs. control (n = 4/group). Two-tailed Student’s t-test [t_6_ = 1.053]. ns, p = 0.332. (**P**) FKPM values for Zbtb7a in astrocyte specific CSDS TRAP-seq data set [GSE139684], with n = 3 control, n = 5 stress: susceptible, n = 4 stress-resilient. 1-way ANOVA [F_2,9_ = 10.01], followed by Tukey’s MC test: control vs. stress susceptible *p = 0.012, stress susceptible vs. stress resilient *p = 0.01, control vs. stress resilient ns, p = 0.989. All data graphed as means ± SEM.

To confirm that ZBTB7A is differentially bound to chromatin in MDD OFC, we carried out footprinting analysis^42^ and calculated ZBTB7A binding predictions within all identified OCR sets. Consistent with motif enrichment analysis, we observed 43.8% more bound ZBTB7A sites that were specific to non-neuronal vs. neuronal cells, with higher occupancy (3.4x) of ZBTB7A sites in non-neuronal MDD cases compared to controls, making it one of the top five most differentially bound TFs genome-wide between MDD and controls (**Figure 2G-H**). One illustrative example demonstrating ZBTB7A binding and increased chromatin accessibility in MDD is *PRR5L* (proline rich 5 like gene), a previously identified MDD biomarker gene involved in stress responsiveness^43, 44^, which displayed increased chromatin accessibility in MDD cases in multiple intronic OCRs. However, the only FDR-significant MDD-specific OCR associated with this gene overlapped with two ZBTB7A binding sites (**Figure 2I**), both of which displayed differential binding in MDD vs. controls.

ZBTB7A was recently identified a master transcriptional regulator of astrocyte inflammatory activation in a recent report using CRISPR screens in human IPSC-derived cells, alongside other more well-characterized factors such as STAT3 and RELA^45^. Consistent with its potential role as a regulator of astrocyte reactivity, pathway analyses revealed that both astrocyte-specific genes and ZBTB7A targets were enriched in differentially expressed genes (DEGs) between MDD vs. controls and in gene targets of non-neuronal promoter OCRs (which were enriched for MDD-related genetic variants) (**Figure 2J, Figure S2C**). Similarly, a previously published MDD OFC RNA-seq dataset also observed ZBTB7A upregulation in MDD, and DEGs in this study were significantly enriched for ZBTB7A targets, demonstrating that altered ZBTB7A regulation in OFC is observed across heterogeneous human MDD cohorts^17^ (**Figure S2D**). In addition, ZBTB7A target genes identified in previously published ChIP-seq datasets showed robust overlap with astrocyte-specific genes (using ARCHS4 human tissue expression reference genes), further suggesting that ZBTB7A is involved in astrocyte function (**Figure S2E**). Consistent with these findings, MDD-specific CREs were found to display significant enrichment for astrocyte/microglia regulatory elements when overlapped with reference panels from human cell-type-specific ATAC-seq data^23^ (note that these two cell-types were sorted together in this dataset) (**Figure 1F**).

Given that we observed increases in both ZBTB7A expression and regulatory activity in human MDD OFC, we next focused on determining whether ZBTB7A expression may also be increased in the context of chronic social stress in mice, an etiologically relevant preclinical model for the study of human MDD. Chronic social defeat stress (CSDS) involves 10 days of exposure to a larger, aggressive mouse during daily social defeat sessions that involve 5-10 minute bouts of physical aggression, followed by 24 hours of sensory exposure to produce continuous psychological stress. CSDS induces robust behavioral deficits in mice that are similar to that observed in human MDD, including reward insensitivity and social avoidance^46^ (**Figure 2K**). Importantly, this paradigm also models natural variation in stress vulnerability, as approximately 30% of wild-type mice that go through CSDS do not exhibit behavioral deficits related to chronic stress and are termed stress-resilient (vs. stress-susceptible). Using this CSDS procedure, we found that Zbtb7a protein was upregulated in bulk OFC tissues from stress-susceptible subjects, but not in control or stress-resilient animals, 48 hours after the final defeat session (*n* = 8 control, *n* = 11 stress-susceptible, *n* = 9 stress-resilient) (**Fig 2L**). We also observed that Zbtb7a expression was persistently increased in OFC of a separate cohort of stress-susceptible mice 21 days after CSDS (*n* = 12 control, *n* = 13 stress-susceptible), demonstrating that Zbtb7a upregulation is maintained for prolonged periods following stress exposures (**Figure S2F-G**). To next determine whether chronic stress leads to *Zbtb7a* upregulation within specific brain cell-types, we utilized Magnetically Activated Cell Sorting (MACs) to isolate astrocyte-, neuron-, and microglia-enriched cell fractions from the OFC of a separate CSDS cohort (*n* = 4/group, with 3 pooled animals/*n*) (**Figure S2H-J**). While we observed ∼2.7x higher expression of *Zbtb7a* mRNA in neurons vs. astrocytes in unstressed animals (consistent with previously published single-cell seq profiles^47^), IHC immunostaining showed that Zbtb7a protein was also expressed at robust levels in mouse OFC astrocytes (**Figure S2K-M**). Importantly, we found that *Zbtb7a* mRNA was increased exclusively in astrocytes following chronic stress exposures, with no significant differences observed in neurons or microglia (**Figure 2M-O**). Finally, we examined *Zbtb7a* expression in an astrocyte-specific Translating Ribosome Affinity Purification coupled to sequencing (TRAP-Seq) dataset (*n* = 3 control, *n* = 5 stress-susceptible, *n* = 4 stress-resilient). Here, we found that *Zbtb7a* mRNA translation was significantly upregulated in frontal cortical (but not hippocampal or striatal) astrocytes of stress-susceptible mice, compared to both control and stress-resilient animals – suggesting that astrocytic *Zbtb7a* levels may correlate with behavioral stress responsivity (**Figure 2P**).

To next determine if ZBTB7A regulates gene targets associated with MDD-specific CREs, we assessed the impact of overexpressing ZBTB7A (OE) in human primary cortical astrocytes using a lentivirus. In this astrocyte-enriched human cell culture system, ZBTB7A OE was found to significantly alter the expression of numerous genes regulated by MDD-specific CREs. ZBTB7A OE also increased the expression of prominent genes within the NF-kB pathway, which was found to be altered in our human MDD dataset (**Figure S2N-Q**). To explore if ZBTB7A might increase in astrocytes under inflammatory conditions, we next treated both human and mouse primary astrocyte-enriched cell cultures with lipopolysaccharide (LPS) to stimulate an inflammatory response, which resulted in a significant upregulation of *ZBTB7A/Zbtb7a* expression compared to saline, further linking this chromatin regulator to cellular reactivity pathways in astrocytes (**Figure S2R-S**).

### Astrocytic Zbtb7a in rodent OFC is necessary to induce behavioral deficits associated with chronic stress

Our human data identified ZBTB7A as an enriched chromatin regulator in glial MDD-specific CREs in OFC, and further validations in mouse models suggested that astrocyte-specific Zbtb7a activity may play important roles in behavioral stress responsivity. Therefore, we next set out to determine whether astrocyte-specific knockdown (KD) of Zbtb7a might be sufficient to attenuate maladaptive behavioral responses to chronic stress. To do so, we designed a *Zbtb7a*-targeting microRNA (miR) construct with a GFP reporter and cloned this construct – vs. a non-gene targeting scrambled miR-GFP control – into an astrocyte-specific GFAP promoter-driven AAV vector for viral packaging into AAV6^38^. We confirmed preferential expression of GFP transgene expression in GFAP+ cells using MACs isolated astrocytes, and validated the efficiency of *Zbtb7a* KD in transduced mouse OFC tissues (**Figure S3A-C**). We next transduced OFC of male mice with AAV6-GFAP-Zbt-miR (Zbt-KD) vs. miRNA-negative-GFP (GFP) viruses prior to CSDS, with half of each viral group being assigned to either control or CSDS conditions (*n* = 7 GFP control, *n* = 9 Zbtb7a KD control, *n* = 18 GFP chronic stress, *n* = 19 Zbtb7a KD chronic stress) (**Figure 3A**). Post-CSDS, we found that astrocyte-specific Zbtb7a KD in OFC was sufficient to fully rescue chronic stress-induced social avoidance observed in GFP-expressing animals, with no significant changes observed in Zbt-KD non-stressed mice (**Figure 3B**). Importantly, Zbtb7a KD also rescued anhedonia-like behavior post-CSDS in two different measures of saccharin reward sensitivity: a Pavlovian cue-reward association task, in which mice learn to associate a signal light with reward delivery, as well as an operant reward learning task requiring lever-pressing in response to a cue light to receive rewards (in a separate cohort of *n* = 7 GFP control, *n* = 7 Zbtb7a KD control, *n* = 8 GFP chronic stress, *n* = 9 Zbtb7a KD chronic stress) (**Figure 3C-F**). Whereas chronically stressed mice learned the reward contingencies of these tasks slower than controls, we observed a significant increase in the number of rewards earned for the Zbtb7a KD chronically stressed mice compared to GFP stressed animals. Together, these results indicate that Zbtb7a activity in OFC astrocytes is a key contributor to behavioral stress responsivity, including social avoidance and reward insensitivity, following chronic psychosocial stress experiences.

**Fig. 3.**
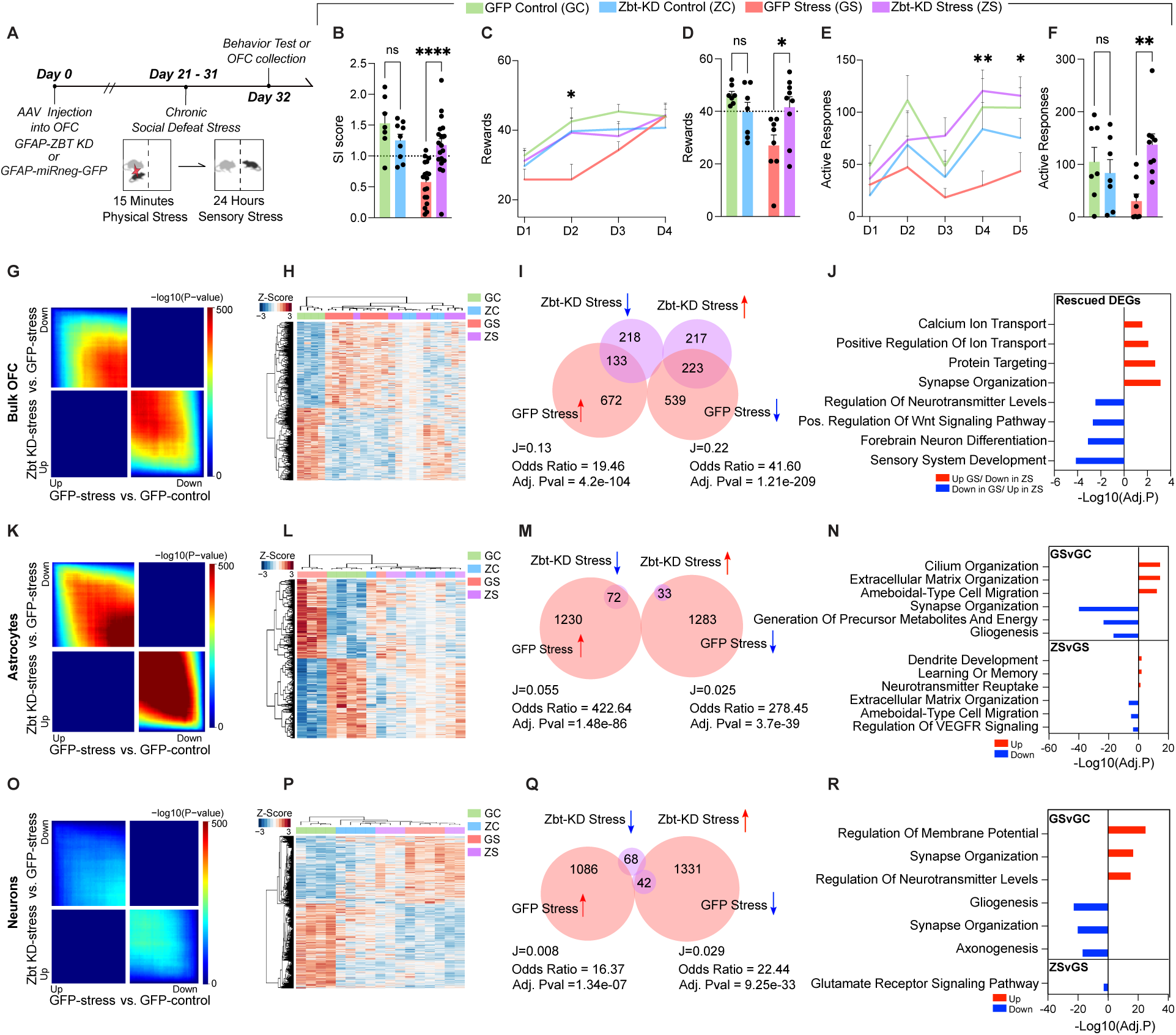
Zbtb7a in rodent OFC astrocytes is necessary to promote chronic stress-induced alterations in behavior and gene expression. (**A**) Schematic of experimental timeline with CSDS paradigm performed after rAAV6 injection into OFC, followed by behavioral test and tissue collection for molecular analyses. (**B**) Social interaction scores. 2-way ANOVA main effect of interaction [F_1,49_ = 13.97], ***p = 0.0005. Sidak’s MC test, GFP control vs. GFP stress ****p< 0.0001. GFP Stress vs. Zbt-KD stress **** p<.0001. Zbt-KD control vs. Zbt-KD stress ns, p=.8663. GFP control vs. Zbt-KD control ns, p = 0.2958. (**C**) Pavlovian cue-reward association task. “D” = Day of task. Mixed Effects analysis, main effect of Test Day x Stress [F_3,83_ = 3.460] *p = 0.0200. (**D**) Individual values for Day two of task shown in (C). 2-way ANOVA main effect of Interaction [F_1,27_ = 8.500] p = 0.0071. Sidak’s MC test GFP control vs. GFP stress **p = 0.0019. GFP stress vs. Zbt-KD stress *p = 0.0119. Zbt-KD control vs. Zbt-KD stress ns, p=.9208. GFP control vs. Zbt-KD control, ns, p = 0.4086. (**E**) Effort-based operant reward learning task on FR1 schedule, “D” = Day of task. Mixed Effects analysis main effect of Virus x Stress [F_1,27_ = 5.835] *p = 0.0228. (**F**) Individual values for Day four of task shown in (E). 2-way ANOVA main effect of Interaction [F_1,27_ = 8.531] *p = 0.0070. Sidak’s MC test GFP control vs. GFP stress, *p = 0.0490. GFP stress vs. Zbt-KD stress **p = 0.0023. Zbt-KD control vs. Zbt-KD stress ns, p=.1759. GFP control vs. Zbt-KD control ns, p = 0.7740. (**G**) RRHO comparing gene expression for the indicated comparisons in bulk OFC tissue. Each pixel represents the overlap between differential transcriptomes, with the significance of overlap of a hypergeometric test color-coded. (**H**) Clustering of groups at 1,583 DE genes (FDR < 0.1) between GFP stress and GFP control in bulk OFC. (**I**) Scaled Venn-diagram and odds ratio test of the overlap between differentially expressed (DE) genes in bulk OFC tissues comparing Zbt-KD stress vs. GFP stress, with GFP stress vs. GFP control. “*J*” indicates the Jaccard index. (**J**) GO analysis for rescued genes in Zbt-KD stress vs. GFP-stress. (**K**) RRHO comparing gene expression for the indicated comparisons in MACS-isolated astrocytes. Each pixel represents the overlap between differential transcriptomes, with the significance of overlap of a hypergeometric test color-coded. (**L**) Clustering of groups at 2,673 DE genes (FDR < 0.1) between GFP stress and GFP control in MACS-isolated astrocytes. (**M**) Scaled Venn-diagram and odds ratio test of the overlap between DE genes in MACS-isolated astrocytes comparing Zbt-KD stress vs. GFP stress, with GFP stress vs. GFP control. “*J*” indicates the Jaccard index. (**N**) GO analysis for gene DEGs in GFP-stress vs. GFP control and Zbt-KD stress vs. GFP-stress, separated by up/down regulation. (**O**) RRHO comparing gene expression for the indicated comparisons in MACS-isolated neurons. Each pixel represents the overlap between differential transcriptomes, with the significance of overlap of a hypergeometric test color-coded. (**P**) Clustering of groups at 2,540 DE genes (FDR < 0.1) between GFP stress and GFP control in MACS-isolated neurons. (**Q**) Scaled Venn-diagram and odds ratio test of the overlap between DE genes in MACS-isolated neurons comparing Zbt-KD stress vs. GFP stress, with GFP stress vs. GFP control. “*J*” indicates the Jaccard index. (**R**) GO analysis for gene DEGs in GFP-stress vs. GFP control and Zbt-KD stress vs. GFP-stress, separated by up/down regulation. All data graphed as means ± SEM.

### Knockdown of Zbtb7a in rodent OFC astrocytes significantly reverses cell-type specific gene expression signatures associated with chronic stress

To explore the molecular changes associated with Zbtb7a KD in the context of chronic stress, we next performed bulk RNA-seq profiling on virally transduced OFC tissues from a separate cohort of animals (*n* = 4 GFP control, *n* = 4 Zbtb7a KD control, *n* = 8 GFP chronic stress, *n* = 8 Zbtb7a KD chronic stress). Threshold-free Rank-Rank Hypergeometric Overlap (RRHO)^48^ analysis revealed transcriptome-wide patterns of reversed gene expression between Zbt-KD stress vs. GFP stress and GFP stress vs. GFP controls, demonstrating that Zbtb7a KD reverses overall gene signatures induced by chronic stress in OFC, maintaining a profile more similar to that of control animals (**Figure 3G, Figure S3E**). Consistent with previous reports that increased Zbtb7a drives NF-kB activation, transcriptome-wide GSEA analysis demonstrated that Zbtb7a KD reversed the upregulation of inflammatory response gene sets induced by chronic stress, with cytokine production being the most significantly downregulated gene set between Zbt-KD stress and GFP stress (**Figure S3F**). Unsupervised clustering of 1,583 DEGs at FDR <.1 between CSDS and controls showed that both Zbt-KD stress and Zbt-KD controls display an intermediate gene expression phenotype that clusters between controls and chronic stress (**Figure 3H**). Odds ratio analysis revealed significant overlap between DEGs that were up in GFP stress and down in Zbt-KD stress (37.8% reversed, Adj.pval = 4.2e-104), and genes that were down in GFP stress and up in Zbt-KD stress (50.5% reversed, Adj.pval = 1.2e-209) (**Figure 3I**). Interestingly, rescued DEGs were also enriched for genes involved in synaptic organization, neurotransmitter regulation, and calcium/ionic transport, suggesting that astrocytic Zbtb7a KD in the context of chronic stress may alter astrocyte function to have cell non-autonomous effects on neuronal transmission (**Figure 3J**).

Therefore, to better define the effects of astrocytic Zbtb7a KD specifically on astrocyte gene expression, we next performed RNA-seq on MACs-isolated astrocytes from Zbtb7a KD vs. GFP groups (+/-) CSDS (*n*=4 GFP control, *n*=4 Zbtb7a KD control, *n*=4 GFP chronic stress, *n*=5, Zbtb7a KD chronic stress, with 3 pooled OFC astrocyte fractions per *n*). Similar to bulk OFC tissues, Zbtb7a KD significantly reversed transcriptome-wide gene expression in astrocytes compared to GFP stress animals (**Figure 3K-L**). Furthermore, approximately 96% (112/117) of the DEGs (FDR<.1) between the Zbtb7a KD stress and GFP stress groups were rescued, including a gene previously identified to be important for chronic stress behavioral responses - *Dusp6*^17^, and the glial-specific glutamate transporter *Slc1a2* (**Figure 3M**). Pathway analysis demonstrated that upregulated pathways in chronic stress were associated with astrocyte reactivity (e.g., cell motility and morphological remodeling), while downregulated genes were involved in critical astrocyte functions, such as metabolic homeostasis and regulation of ionic transport and synaptic signaling — pathways that were also enriched for DEGs that were rescued by Zbtb7a KD (**Figure 3N**).

To examine potential cell non-autonomous effects of astrocytic Zbtb7a KD, we next profiled MACs-isolated OFC neurons from the same cohort of Zbtb7a KD vs. GFP animals (+/-CSDS) (*n* = 4 GFP control, *n* = 4 Zbt-KD control, *n* = 4 GFP chronic stress, *n* = 5, Zbt-KD chronic stress, with 3 pooled OFC neuronal fractions per *n*) (**Figure 3O-P**). Comparing these data with astrocyte-specific profiles, we confirmed that both astrocyte and neuronal fractions demonstrated cell-type specific expression patterns for respective population markers^49^ (**Figure S3H**). Interestingly, we found that Zbtb7a KD specifically within astrocytes also led to cell non-autonomous effects on neuronal gene expression in the context of stress, including a reversal of genes associated with glutamate transmission (**Figure 3Q**). These data suggest that during chronic stress, astrocytes lose normal homeostatic processes that may have downstream consequences on OFC neuronal activity, effects that are attenuated by reducing Zbtb7a activity in astrocytes specifically.

### Zbtb7a regulates chromatin accessibility patterns in astrocytes associated with chronic stress

Since ZBTB7A was previously identified as a chromatin remodeling protein involved in multiple cell-signaling pathways, including NF-kB inflammation^40^, we next sought to confirm whether astrocyte-specific manipulations of Zbtb7a alter chromatin accessibility patterns in the context of chronic stress. To do so, we performed ATAC-seq on MACs-isolated astrocytes from virally-infected OFC tissues from the four groups of animals described above (*n* = 4 GFP control, *n* = 4 Zbt-KD control, *n* = 5 GFP stress, *n* = 5 Zbt-KD stress, with each *n* composed of 3 pooled OFC samples), and found that promoters displaying chronic stress-induced accessibility were enriched for Zbtb7a targets, as were less accessible promoter regions in Zbt-KD stress vs. GFP stress animals, indicating that these chromatin profiles reflect Zbtb7a regulatory activity in astrocytes (**Figure S3N**). Differential accessibility analyses demonstrated that Zbtb7a KD rescued astrocyte-specific chromatin accessibility patterns induced by chronic stress, with 42.3% (603/1391, Adj. Pval = 2e-89) of up events and 65.5% (2044/3117, Adj.Pval = 0e+00) of down events displaying opposing accessibility compared to GFP stress mice (**Figure S3P**). In addition, Zbt-KD stress ATAC-seq profiles correlated significantly with Zbt-KD stress gene expression changes detected in our astrocyte-specific RNA-seq dataset, and exhibited a reversed pattern of anti-correlation with chronic stress OCRs (**Figure S3Q-R**). Rescued OCRs were found to be enriched for pathways involved in astrocyte reactivity, including ion homeostasis, ECM alterations, and cellular morphogenesis (**Figure S3S**). Importantly, among the genes reversed by Zbtb7a KD for both gene expression and chromatin accessibility were the astrocyte-specific glutamate clearance transporter gene *Slc1a2* (also known as *Eaat2)*, which modulates neuronal excitability through maintenance of glutamatergic tone (note that *Slc1a2* has consistently been shown to be downregulated following chronic stress^50, 51^). These findings again highlight that Zbtb7a-mediated astrocyte dysfunction during chronic stress may affect neuronal function through regulation of glutamate clearance and synaptic excitability (**Figure S3T**).

### ZBTB7A overexpression in astrocytes is sufficient to induce behavioral deficits following an innocuous mild subthreshold stressor

In order to explore mechanistic roles for astrocytic Zbtb7a upregulation in stress vulnerability and to assess whether increased Zbtb7a is sufficient to elicit a behavioral stress response, we packaged an OE construct for *ZBTB7A* into the same AAV6-GFAP viral vector used for KD experiments (**Figure S4A-E**). Our previous data suggested that increased ZBTB7A may be associated with increased vulnerability to stress-related behavioral deficits. Therefore, for OE experiments, we utilized the sub-threshold social defeat paradigm (SSDS) – which involves only a single day of social defeat and does not induce behavioral deficits in wild-type animals^52^ -- to assess whether ZBTB7A OE in OFC astrocytes is sufficient to promote a pro-susceptibility behavioral phenotype after a normally innocuous mild stressor (*n*=10 GFP control, *n*=8 ZBT-OE control, *n*=20 GFP SSDS, *n*=18 ZBT-OE SSDS) (**Figure 4A**).

**Fig. 4.**
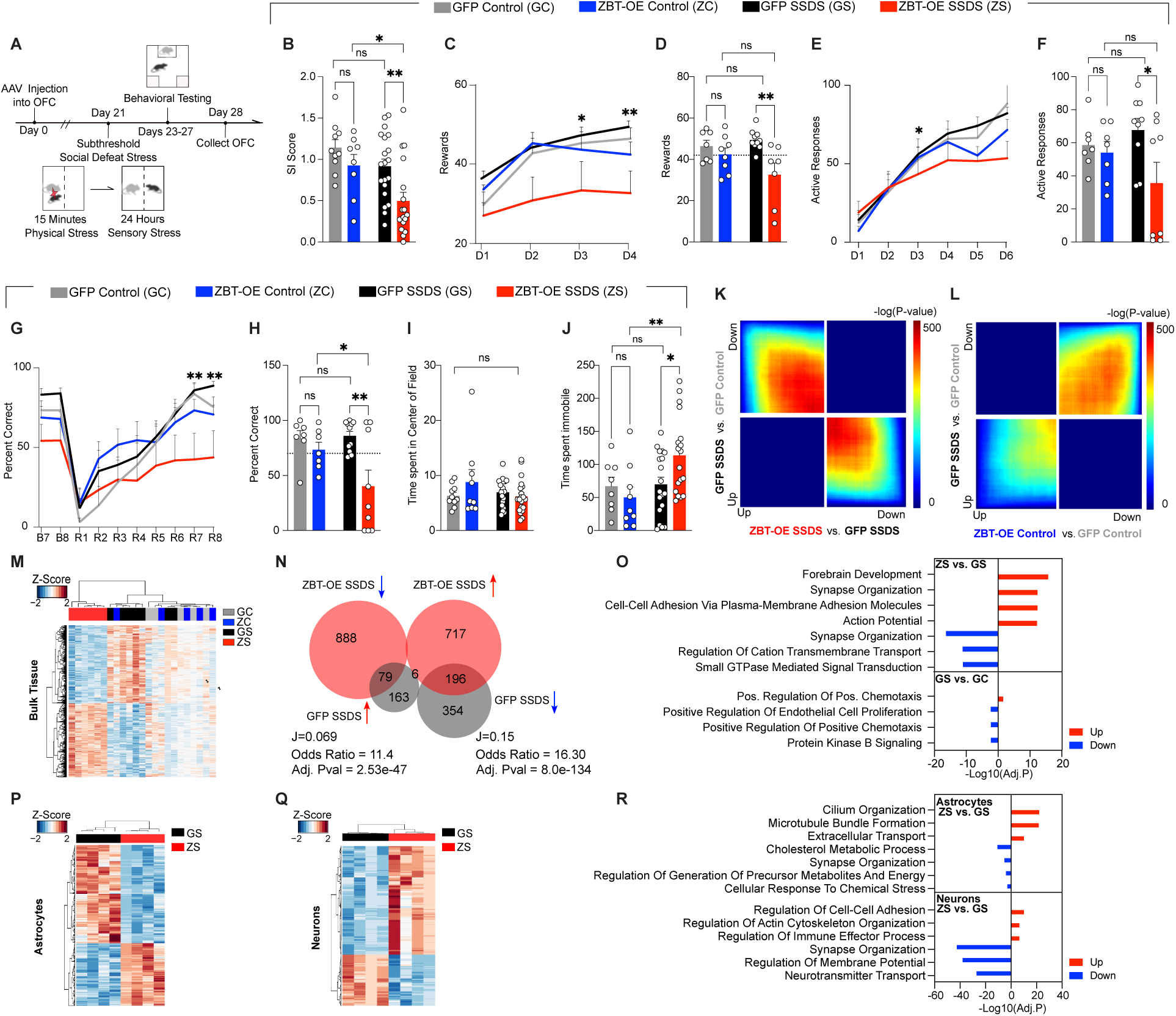
ZBTB7A in mouse OFC astrocytes is sufficient to induce chronic stress-mediated alterations in chromatin accessibility, gene expression, and behavior. (A) Schematic of experimental timeline with subthreshold SSDS mild stress paradigm performed after rAAV6 injection into OFC, followed by behavioral tests and tissue collection for RNA-seq. (B) Social interaction. 2-way ANOVA main effect of stress [F_1,52_ = 8.144], **p = 0.0062, main effect of virus [F_1,52_ = 7.730], **p = 0.0075. Sidak’s MC test, GFP control vs. GFP SSDS ns, p = 0.2788. GFP SSDS vs. ZBT-OE SSDS **p = .0041. ZBT-OE control vs. ZBT-OE SSDS *p = .0286. GFP control vs. ZBT-OE control n.s. p = .4480. (**C**) Pavlovian cue-reward association task. “D” indicates Day of test. 3-way ANOVA, main effect of Virus x Stress [F_1,29_ = 5.291] *p = 0.0288. (**D**) Individual values for day 2 of task shown in (C). 2-way ANOVA, main effect of virus [F_1,28_ = 9.759], p = **0.0041. Sidak’s MC test, GFP control vs. GFP SSDS ns, p=0.651. GFP SSDS vs. ZBT-OE SSDS **p = .0021. ZBT-OE control vs. ZBT-OE SSDS ns, p = 0.146. (**E**) Operant reward task, FR1. “D” indicates Day of test. 3-way ANOVA, main effect of Test Day x Virus [F_5,149_ = 2.823] *p = 0.0182. (**F**) Individual values for day 3 of task shown in (E). 2-way ANOVA, main effect of virus [F_1,27_ = 4.408], *p = 0.0453. Sidak’s MC test, GFP control vs. GFP SSDS ns, p=0.709. GFP SSDS vs. ZBT-OE SSDS *p = .0218. ZBT-OE control vs. ZBT-OE SSDS ns, p = 0.282. (**G**) Percent correct trials in reversal learning paradigm. “B” indicates Baseline day, “R” indicates Reversal phase day. 3-way ANOVA, main effect of Test day x Virus [F_9,261_ = 4.529] p < 0.0001. (**H**) Individual values for day 7 as shown in (G). 2-way ANOVA, main effect of virus [F_1,30_ = 9.017], **p = 0.0054. Sidak’s MC test, GFP control vs. GFP SSDS ns, p=0.9797. GFP SSDS vs. ZBT-OE SSDS **p = .0013. ZBT-OE control vs. ZBT-OE SSDS *p = 0.0389. GFP control vs. ZBT-OE control n.s., p = 0.7280. (**I**) Time spent (s) in the center of the field during open field test. 2-way ANOVA ns, (**J**) Forced Swim tests. 2-way ANOVA main effect of interaction [F_1,50_ = 4.129], *p = 0.0475, main effect of stress [F_1,50_ = 4.993], *p = 0.0475. Sidak’s MC test, GFP control vs. GFP SSDS ns, p=0.9876. GFP SSDS vs. ZBT-OE SSDS **p = 0.0070. (**K-L**) RRHO comparing gene expression between indicated comparisons, in the context of mild stress. (**M**) Clustering at 1,929 DE genes between ZBT-OE SSDS and GFP SSDS. (**N**) Scaled Venn-diagram and odds ratio test of the overlap between differentially expressed (DE) genes in bulk OFC tissues comparing ZBT-OE stress vs. GFP SSDS, with GFP SSDS vs. GFP control. “*J*” indicates the Jaccard index. Note for GFP SSDS vs. GFP control, DEGs were defined at pval < 0.05 (**O**) GO analysis for gene DEGs in ZBT-OE SSDS vs. GFPSSDS and GFP SSDS vs. GFP control, separated by up/down regulation. All data graphed as means ± SEM. (**P**) Clustering at 715 DE genes between ZBT-OE SSDS and GFP SSDS astrocytes (n = 4/group). (**Q**) Clustering at 1,191 DE genes between ZBT-OE SSDS and GFP SSDS neurons (n = 4/group). (**R**) GO analysis for DE genes (FDR < 0.1) between ZBT-OE SSDS and GFP SSDS groups in MACs-isolated astrocytes and neurons, separated by up/down regulation.

We found that astrocyte-specific ZBTB7A OE significantly increased behavioral deficits following acute stress compared to GFP mice, including heightened social avoidance and anhedonic reward insensitivity in the saccharin Pavlovian and operant reward tasks (in a separate cohort of mice, *n* = 7-10/group) (**Figure 4B-F**). ZBTB7A OE + acute stress also induced deficits in reward-based reversal learning, which is a well characterized OFC-dependent task^53^, suggesting that ZBT-OE in astrocytes impairs OFC function following a mild stressor (**Figure 4G-H**). Furthermore, although ZBT-OE SSDS animals did not display significant differences in anxiety-like behaviors in the open field test, they did exhibit a significant increase in immobility in the forced swim test (**Figure 4I-J**). In contrast, GFP-SSDS mice displayed distinct proadaptive behaviors in response to acute stress, with no significant differences observed between GFP SSDS mice and GFP controls. Notably, ZBTB7A OE alone did not affect stress-related behaviors in ZBT-OE control mice. This is in agreement with previous reports that ZBTB7A acts mainly to transduce cellular signals through orchestration of chromatin accessibility^40^, and in the absence of NF-kB activation, ZBTB7A OE does not induce an inflammatory response on its own – though it may prime chromatin states toward heightened stress responses following subsequent adverse experiences^40^.

### ZBTB7A overexpression in astrocytes induces transcriptome-wide alterations in gene expression and chromatin accessibility related to inflammatory signaling and neuroactive communication following a mild stress

To explore the molecular correlates of these behavioral results, we next performed bulk RNA-seq on virally transduced OFC tissues -/+ SSDS (*n* = 5 GFP control, *n* = 5 ZBT-OE control, *n* = 7 GFP SSDS, *n* = 6 ZBT-OE SSDS, 1 OFC per *n*). Both RRHO analysis and unsupervised clustering of DEGs (1,929 genes, FDR<.1) between the two SSDS groups revealed a robust pattern of transcription induced by ZBTB7A OE, while the ZBT-OE control exhibited a positive correlation with GFP SSDS – indicating that ZBTB7A OE in the absence of a mild stressor does not disrupt overall transcriptomic states, in agreement with our behavioral data (**Figure 4K-M**).

In addition, although there were only a small number of DEGs between GFP control and GFP SSDS (19 genes, FDR<.1), they included well-characterized stress-related genes, such as an increase in the resilience-related gene *Fkbp5* and a decrease in the inflammatory cytokine *Cxcl12*. Transcriptome-wide, inflammatory gene sets were found to be significantly downregulated in GFP SSDS vs. GFP control mice, demonstrating that the behavioral resilience observed after an exposure to acute stress in GFP SSDS mice likely involves pro-adaptive transcriptional responses (**Figure S4F**). Overlaps between significant DEGs (FDR < 0.1) in ZBT-OE SSDS vs. GFP SSDS and DEGs with a more relaxed cutoff (Pval <.05) in GFP SSDS vs. GFP Control revealed significant reversal of genes in both directions (32.6%, Adj. Pval = 2.53e-47 and 35.6%, Adj. Pval = 8.0e-134), suggesting that overexpression of ZBTB7A in astrocytes reverses proadaptive gene expression responses associated with resilience to a mild stressor (**Figure 4N**). ZBT-OE + mild stress altered molecular pathways related to astrocyte activation, such as regulation of ionic transport, cellular adhesion/chemotaxis, and synaptic organization, all of which were oppositely regulated in the GFP-SSDS vs. GFP control group (**Figure 4O**). Finally, flow cytometry analysis confirmed that astrocytic ZBTB7A OE + acute stress significantly increased functional markers of neuroinflammation in OFC, with higher percentages of microglia expressing activated markers observed (note that the overall number of astrocytes or microglia was not altered between conditions) (**Figure S4G-H**).

Next, we focused on differences between the two acute stress groups, where ATAC-seq and RNA-seq profiling of MACs-isolated OFC astrocytes from a separate cohort of mice (*n* = 4 GFP SSDS, *n* = 4 ZBT-OE SSDS) revealed robust changes in accessibility (6,094 differentially accessible regions) that correlated significantly with observed differential gene expression patterns (715 DE genes, FDR<.1) (**Figure S4I-K**). Importantly, astrocyte-specific ZBTB7A OE + mild stress induced chromatin accessibility changes that overlapped significantly with those detected following chronic stress (Adj. Pval = 4e-152 for downregulated genes, Adj.Pval = 5e-208 for upregulated genes), suggesting that ZBTB7A-mediated chromatin remodeling may be a central regulatory feature controlling astrocytic dysfunction during chronic stress exposures (**Figure S4L-M**). Indeed, ZBTB7A OE + acute stress induced similar alterations in gene expression to those observed following chronic stress and were associated with astrocyte reactivity pathways, including cellular morphology and synaptic regulation (**Figure 4P-R**). In addition, neuronal specific RNA-seq profiles demonstrated an increase in cellular stress response, as well as decreases in neurotransmitter transport and synaptic organization, indicating that astrocyte-specific ZBTB7A OE may induce behavioral stress susceptibility through modulation of astrocytic-induced OFC neuronal hyperactivity (**Figure 4Q-R**).

### ZBTB7A overexpression in OFC astrocytes potentiates synaptic transmission

To determine whether ZBTB7A upregulation is associated with changes in astrocyte calcium signaling, we next imaged 2D mouse primary mixed cortical cultures of neurons and glia (including astrocytes) using the genetically encoded calcium indicator GCaMP6f. Primary mixed cultures were transduced with AAV1-hSyn-GCaMP6f or AAV5-gfaABC1D-cyto-GCaMP6fto to ensure selective expression solely in neurons or astrocytes, respectively (**Figure S5A-E**). To elicit a subthreshold-like adaptive cellular response, we treated cultures with a low-dose of LPS (LPS_low_), approximately 1-10% of a typical inflammatory dose^54–56^, which has previously been used to examine neuroprotective effects of mild LPS treatment in mixed culture models^55, 57–59^. In GCaMP6f astrocytes, ZBT-OE significantly increased calcium transient frequency both at baseline and after LPS_low_ treatment compared to an empty vector control virus, suggesting that ZBTB7A OE disrupts adaptive astrocyte plasticity to subthreshold stressful stimuli (**Figure S5B**). Furthermore, neuronal calcium events, which are a proxy marker for action potentials, were also found to be significantly increased in co-cultures treated with the astrocyte-specific ZBT-OE virus + LPS_low_ compared to empty vector controls (**Figure S5C**). These data indicate that ZBTB7A OE in astrocytes leads to increased astrocyte activity, impairing both astrocytic and neuronal adaptations to a mild stressful stimulus.

To assess if these astrocytic ZBTB7A-induced increases in neuronal activity occur *in vivo*, we next investigated whether astrocyte specific ZBTB7A OE affects functional measures of synaptic transmission in mouse OFC following SSDS. We utilized our GFAP-driven AAV virus to OE either ZBTB7A or GFP in OFC astrocytes (+/-) SSDS, followed by electrophysiological slice recordings to assess the impact of ZBTB7A OE on synaptic transmission post-exposure to a SSDS mild stressor (**Figure 5A**). We first plotted an input-output (I-O) curve of field excitatory postsynaptic potentials (fEPSPs) in response to presynaptic stimuli. We observed a significant increase in I-O curves in ZBT-OE SSDS vs. GFP SSDS, suggesting that ZBTB7A OE in astrocytes induces potentiation of postsynaptic responses following an acute stress (**Figure 5B-C**). We next applied stimulation protocols to assess the dynamics of presynaptic vesicle mobilization and release. During the stimulation train, we detected a significant difference in fEPSP amplitude between ZBT-OE SSDS vs. GFP SSDS, suggesting a faster depletion of the readily releasable pool of vesicles, which is correlated with higher probability of presynaptic release (**Figure 5D-E**)^60^. Together, these data indicate that astrocytic ZBTB7A OE + acute stress is sufficient to induce increased OFC neuronal excitability, a hallmark feature of human MDD.

**Fig. 5.**
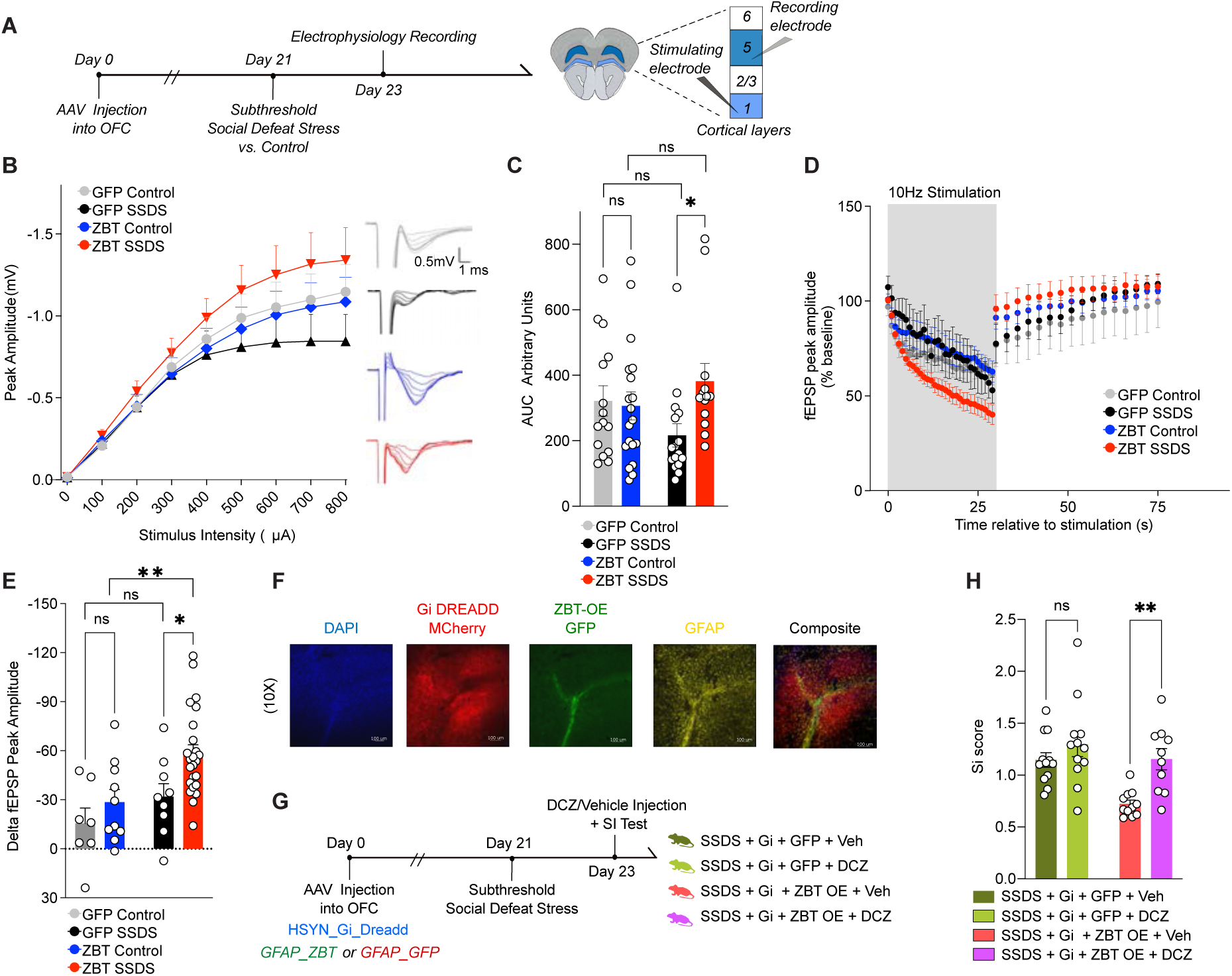
ZBTB7A in mouse OFC astrocytes induces cell non-autonomous neuronal hyperexcitability to mediate stress susceptibility. (**A**) Schematic of experimental timeline with subthreshold stress paradigm performed after AAV6 injection into OFC, followed by slice electrophysiology recordings. (**B**) Input-output (I-O) curve constructed by recording fEPSPs in response to stimuli ranging from 100-800 µA. 3-way ANOVA, main effect of Stimulus Intensity x Virus x Stress [F_8,480_ = 2.626] **p = 0.0080. (**C**) Individual values for (I-O) curve, area under curve (A.U.C). 2-way ANOVA main effect of Interaction [F_1,59_ = 4.062], *p = 0.0484. Sidak’s MC test, GFP control vs. GFP stress ns, p=0.1923. GFP Stress vs. ZBT-OE stress *p = 0.0295. ZBT-OE control vs. ZBT-OE stress n.s., p = 0.4230. GFP control vs. ZBT-OE control n.s., p= 0.9597. (**D**) Rundown stimulation from a single 30-s train delivered at 10 Hz. The percentage change in fEPSP amplitude from baseline was calculated during and post-10Hz stimulation. 3-way ANOVA, main effect of Stimulus x Virus [F_29,1334_ = 3.376] ***p < 0.0001, main effect of stress x virus [F_1,46_ = 4.356] *p = 0.0425. (**E**) Individual values for delta fEPSP amplitude (% baseline) between end of 10Hz stimulation and 1s after end of stimulation train. 2-way ANOVA main effect of virus [F_1,46_ = 6.115], p = 0.0172, main effect of stress [F_1,46_ = 8.454], **p = 0.0056. Sidak’s MC test, GFP control vs. GFP stress ns, p=0.5172. GFP Stress vs. ZBT-OE stress *p = 0.0207. ZBT-OE control vs. ZBT-OE stress **p = 0.0059. GFP control vs. ZBT-OE control n.s., p= 0.5172. (**F**) IHC validation of hsyn-hM4D(Gi)-mCherry (in red) and GFAP-ZBT OE (in green) localized in astrocytes (GFAP, in yellow) and DAPI (in blue). Images taken at 10x magnification. (**G**) Experimental scheme of chemogenetics experiment, in which SSDS is performed on a cohort of mice expressing hM4D(Gi)-mCherry (+/-) ZBT OE, (+/-) DCZ. (**H**) Social interaction. 2-way ANOVA main effect of Virus [F_1,41_ = 10.11], **p = 0.0028, main effect of agonist [F_1,41_ = 10.65], **p = 0.0022. Sidak’s MC test, Gi + GFP stress + vehicle vs. Gi + GFP stress + DCZ ns, p=0.3880. Gi + ZBT-OE stress + vehicle vs. Gi + ZBT-OE stress + DCZ **p = 0.0040. All data graphed as means ± SEM.

### Astrocytic ZBTB7A-induced OFC neuronal hyperexcitability mediates behavioral vulnerability to stress

To determine if increased OFC neuronal activity represents a functional link between ZBTB7A-mediated astrocyte reactivity and behavioral vulnerability to stress, we utilized an inhibitory Designer Receptors Exclusively Activated by Designer Drugs (DREADD)-based chemogenetic approach to silence OFC neurons while simultaneously overexpressing ZBTB7A in surrounding astrocytes. We first confirmed that I.P. injection of the DREADD agonist Deschloroclozapine^61^ (DCZ) on its own did not alter previously observed patterns of behavioral deficits in ZBT OE vs. GFP mice post-stress in the absence of the DREADD, and that neuronal G_i_ DREADD expression without DCZ agonist activation does not affect previously observed SI phenotypes (**Figure S5F-H**). We then performed the SSDS paradigm on a separate cohort of male mice that were injected intra-OFC with the both the pAAV-hSyn-hM4D(G_i_)-mCherry vector to express the inhibitory G_i_ DREADD in neurons, and either the AAV-GFAP-*ZBTB7A* OE virus or the rAAV6-GFAP-GFP empty control vector in astrocytes (**Figure 5F**). Note that both groups underwent the SSDS paradigm. Prior to the social interaction test, half of each viral group was injected with either vehicle or DCZ to silence OFC neuronal firing (**Figure 5G**). In the G_i_ + vehicle-injected mice, ZBT OE + acute stress resulted in reduced social interaction behavior vs. GFP SSDS, as previously observed. However, for DCZ-injected mice, we observed a significant rescue in SI deficits, indicating that silencing of OFC activity prior to the social interaction test led to amelioration of the pro-stress susceptibility effects of astrocyte-specific ZBTB7A OE in the context of a mild stressor (**Figure 5H**). Together, these findings point to an astrocytic ZBTB7A-induced increase in synaptic connectivity driving maladaptive stress susceptibility, which is consistent with clinical reports of astrocyte dysfunction and neural hyperactivity in human MDD OFC^10^.

## Discussion

Overall risk for MDD is determined by complex interactions between genetic and environmental factors that disrupt frontolimbic function. Although MDD has primarily been studied in the context of neuronal plasticity, recent studies suggest that dysregulation of glial cell activity, and astrocytes specifically, may be a key contributor to MDD pathophysiology^35, 36, 51, 62^. The OFC exhibits distinct functional changes from other frontal cortical regions in chronic stress and MDD^11^, however the molecular pathways driving these alterations within specific cell-types are not well understood. Here, implementing both RNA-seq and FANs-coupled ATAC-seq in postmortem MDD OFC tissues, we identified significant glial dysfunction in depressed individuals vs. controls, including alterations in inflammatory pathways, cellular metabolism, and ionic homeostasis. Using unbiased cell-population-specific epigenomic profiling, we identified a key chromatin regulator of MDD-specific CREs in human depression, ZBTB7A, which was found to be upregulated in MDD and displayed significantly higher motif occupancy in MDD non-neuronal cells vs. controls. We validated the relevance of this chromatin regulator to astrocyte-mediated chronic stress phenotypes using preclinical mouse models for the study of MDD, and demonstrated that astrocyte-specific ZBTB7A regulatory activity in OFC bidirectionally mediates molecular, electrophysiological, and behavioral alterations induced as a consequence of chronic stress exposures.

Our human postmortem ATAC-seq data, together with functional validation experiments in preclinical mouse models, demonstrated that astrocytic ZBTB7A may act as a pathogenic driver of astrocyte dysfunction in MDD by inducing inflammatory reactivity and compromising normal astrocyte-mediated regulation of synaptic function^63^ and glutamatergic signaling^63, 64^. Our mouse data also revealed that ZBTB7A upregulation acts to reverse normal adaptive mechanisms that promote stress resilience. Importantly, our rodent data are consistent with clinical reports of OFC neural hyperactivity in human MDD^10^, and raise the intriguing potential of targeting astrocytic ZBTB7A, as well as its downstream substrates mediating OFC dysfunction, therapeutically. Overall, these findings support a critical role for astrocyte plasticity in the pathophysiology of MDD and stress-related disorders and highlight the power of using epigenomic profiling to investigate novel regulatory mechanisms driving aberrant cellular phenotypes in complex disease states.

## Methods

### Human postmortem samples

Postmortem human orbitofrontal cortex (Brodmann Area 11) tissues from 39 Caucasian subjects (20 cases, 19 controls) were obtained from the Human Brain Collection at the University of Texas Southwestern (UTSW) (IRB approval for tissue banking at UTSW). Tissue preservation was achieved as previously described^65^. Brains were placed on wet ice and transported to the UTSW Brain Bank facilities. Tissues were sliced, flash frozen in 2-methylbutane at −40°C, and stored in sections conserving anatomical landmarks at −80°C. OFC tissues were later sectioned from frozen slices. For each subject, the cause of death was determined by the Coroner Office, and toxicological screens were performed to obtain information on medication and illicit substance use at their time of death. The MDD group consisted of 20 (9 male and 11 female) individuals who met the Structured Clinical Interview for DSM-V (Diagnostic and Statistical Manual of Mental Disorders-V) Axis I Disorders: Clinician Version (SCID-I) criteria for Major Depressive Disorder. The control group comprised 19 subjects (12 male and 7 female Caucasians) with no history of MDD. Groups were matched for age, post-mortem interval and RNA integrity number (RIN). For all subjects, psychological autopsies were performed, giving us access to detailed information on psychiatric and medical histories, as well as other relevant clinical and sociodemographic data (see **Table S1)**.

### FANS sorting of neuronal and non-neuronal nuclei

50mg of frozen brain tissue was homogenized in cold lysis buffer (0.32M Sucrose, 5 mM CaCl2, 3 mM Mg(Ace)2, 0.1 mM, EDTA, 10mM Tris-HCl, pH8, 1 mM DTT, 0.1% Triton X-100) and filtered through a 40µm cell strainer. The flow-through was underlaid with sucrose solution (1.8 M Sucrose, 3 mM Mg(Ace)2, 1 mM DTT, 10 mM Tris-HCl, pH8) and subjected to ultracentrifugation at 24,000 rpm for 1 hour at 4°C. Pellets were thoroughly resuspended in 500µl DPBS and incubated in BSA (final concentration 0.1%) and anti-NeuN antibody (1:1000, Alexa488 conjugated, Millipore) under rotation for 1 hour, at 4 °C, in the dark. Prior to FANS sorting, DAPI (Thermoscientific) was added to a final concentration of 1µg/ml. DAPI positive neuronal (NeuN+) and non-neuronal (NeuN-) nuclei were sorted into tubes pre-coated with 5%BSA using a BD-FACSAria flow cytometer (BD Biosciences) equipped with a 70μm nozzle **(Figure S6)**. 39 tissue dissections from 1 brain region were subjected to FANS, resulting in 78 (39 NeuN- and 39 NeuN+) distinct nuclear populations.

### RNA-sequencing

For human postmortem OFC, ∼25mg of pre-sectioned flash-frozen tissue was utilized for RNA extraction. For mouse studies, animals were euthanized, and brains were removed whole and flash frozen (for bulk sequencing), or processed fresh for cell-type specific isolation with magnetically-activated cell sorting (MACs). Brains were sectioned at 100 µm on a cryostat (bulk) or brain block (MACs), and GFP was illuminated using a NIGHTSEA BlueStar flashlight to microdissect virally infected tissues with a 2mm punch. For both human and mouse experiments, OFC tissues were homogenized in Trizol (Thermo Fisher), and RNA was isolated on RNeasy Minelute Microcolumns (Qiagen) following manufacturer’s instructions. Following elution, samples were enriched for mRNA via polyA tail selection beads, and mRNA libraries were prepared using the Illumina Truseq RNA Library Prep Kit V2 (#RS-122-2001). Libraries were pooled and sequenced on the Illumina Novaseq platform, with an average read count of approximately 20 million paired-end reads per sample. RNA-seq data was pre-processed and analyzed as previously described^66^. Briefly, FastQC (Version 0.72) was performed on the concatenated replicate raw sequencing reads from each library to ensure minimal PCR duplication and sequencing quality. Reads were aligned to the hg38 or mouse mm10 genome using HISAT2 (Version 2.1.0) and annotated against Ensembl v90. Multiple-aligned reads were removed, and remaining transcript reads were counted using featurecounts (Version 2.0.1). For mouse RNA-sequencing experiments with multiple groups, RUVg^67^ was performed to normalize read counts based on empirically determined control genes that do not vary between control and stress groups (i.e. genes with p-val > 0.5 based on a first-pass differential expression analysis performed prior to RUVg normalization). For human RNA-seq and mouse RNA-seq experiments with two groups, RUVr^67^ was performed to normalized read counts based on the residuals from a first-pass GLM regression of the unnormalized counts on the covariates of interest. The number of factors of variation, or RUV *k*, for each experiment is listed in **Table S5**). DESEQ2^68^ (Version 2.11.40.6) was used to perform pairwise differential expression analyses between indicated comparisons. Differentially expressed (DE) genes (listed in **Table S5** for each experiment) were defined at FDR<0.1. Threshold free Rank-Rank Hypergeometric Overlap (RRHO) maps were generated to visualize transcriptome-wide gene expression concordance patterns as previously described^48^, using RRHO2 (Version 1.0). For RRHO comparing ATAC-seq vs. RNA-seq, signed log p-value from the RNA-seq DESEQ2 output was ranked for each transcript that was also associated with a differentially accessible peak in the ATAC-seq.

For the human MDD dataset, we used the WGCNA package (Version 1.71)^22^ to construct the co-expression network for the top 2000 most variable genes in the set. We chose a suitable soft threshold power of 7 for scale-free network construction with the function pickSoftThreshold. The resulting gene co-expression network was visualized as the heatmap based on dissimilarity of TOM with hierarchical clustering dendrogram, and the number of genes in each module was counted. The correlation between modules and the trait of MDD was assessed by the Pearson correlation coefficients, with students *t*-test, and a *p* value of < 0.05 was considered statistically significant. Gene ontology (GO) enrichment analysis was performed for genes in each significant module (and for GO analyses on DE genes from other experiments) with gprofiler(GO), idep (TRANSFAC/JASPAR databases) with total detected genes as background, and enrichR (for cell-type and human disease databases) to test for overrepresented gene categories in our list of DE genes. FDR for representative GO terms from the top 10 terms is calculated based on nominal P-value from the hypergeometric test. Gene Set Enrichment Analysis was performed using the ClusterProfiler package (Version 4.6.0) against GO to calculate gene set enrichment scores, and gene sets were ranked by adj. p-value^69^. Odds Ratio analyses were carried out on DE gene lists using the *GeneOverlap* R package version 1.26.0^70^.

### Generation of human ATAC-seq libraries

ATAC-seq reactions were performed using an established protocol^71^ with minor modifications. Following FANS, 50,000 sorted nuclei were centrifuged at 500 ×g for 10 min, 4°C. Pellets were resuspended in transposase reaction mix (25 μL 2x TD Buffer (Illumina Cat #FC-121-1030) 2.5 μL Tn5 Transposase (Illumina Cat #FC-121-1030) and 22.5 μL Nuclease Free H2O) on ice. Samples were incubated at 37°C for 30 min and then purified using the MinElute Reaction Cleanup kit (Qiagen Cat #28204) according to the manufacturer’s instructions. Following purification, library fragments were amplified using the Nextera index kit (Illumina Cat #FC-121-1011), under the following cycling conditions: 72°C for 5 minutes, 98°C for 30 seconds, followed by thermocycling at 98°C for 10 seconds, 63°C for 30 seconds, and 72°C for 1 minute for a total of 5 cycles. In order to prevent saturation due to over-amplification, a 5µl aliquot was then removed and subjected to qPCR for 20 cycles to calculate the optimal number of cycles needed for the remaining 45 μL reaction. The additional number of cycles was determined as follows: (1) Plot linear Rn vs. Cycle (2) Calculate the # of cycles that corresponds to 1⁄4 of maximum fluorescent intensity. In general, we found adding 4-6 cycles to this estimate yielded optimal ATAC-seq libraries, as determined by analysis on Bioanalyzer High Sensitivity DNA Chips (Agilent technologies Cat#5067-4626). Libraries were amplified for a total of 13–19 cycles. Following PCR, ATAC-seq libraries were resolved on 2% agarose gels and fragments ranging in size from 100bp-1Kbp were excised and purified (Qiagen Minelute Gel Extraction Kit – Qiagen Cat#28604). Libraries were quantified by quantitative PCR (KAPA Biosystems Cat#KK4873) prior to sequencing. Libraries were sequenced on Hi-Seq2500 (Illumina) obtaining 2×50 paired-end reads. After quality controls (see below), 70 ATAC-seq libraries were retained for downstream analysis.

### Data Processing

We provide a summary of the data processing pipeline to the right. The preprocessing of ATAC-seq samples involved the following steps: (1) per sample processing, (2) joint processing for quality control, and (3) analyses of samples meeting the quality control criteria. Yellow: input data. Blue: analyses. Green: processed data.

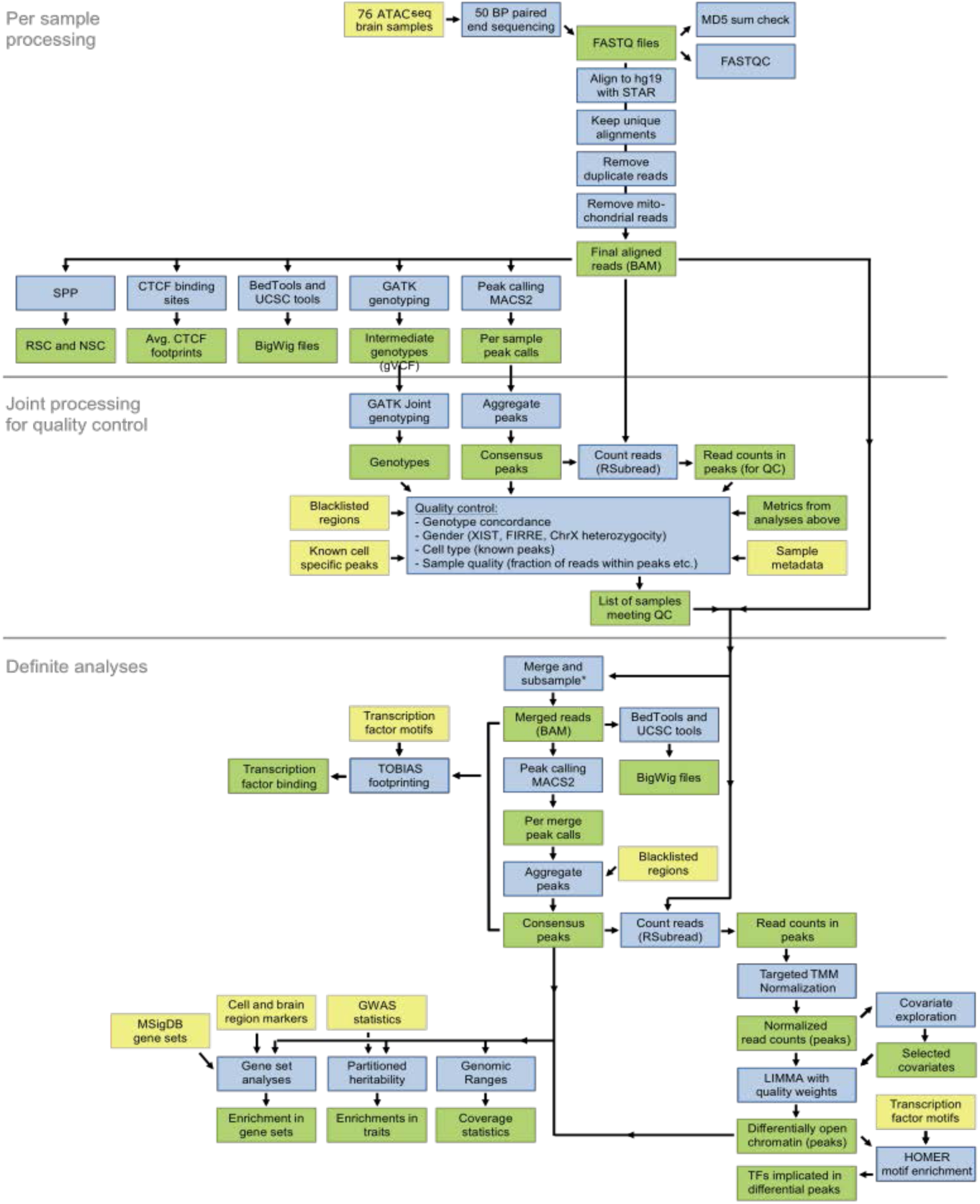

### Alignment

Raw sequencing reads were generated by the sequencing center demuxed and with adaptors trimmed. FASTQ files were linked to the sample clinical and demographics metadata based on pooling ID’s and barcodes. Reads were subsequently aligned to the hg19 reference genome with the pseudoautosomal region masked on chromosome Y with the STAR aligner (v2.5.0)^72^, using the following parameters: *-- alignIntronMax 1, --outFilterMismatchNmax 100, --alignEndsType EndToEnd, -- outFilterScoreMinOverLread 0.3, --outFilterMatchNminOverLread 0.3*. Having a coordinate-sorted BAM, we further excluded reads that: (1) were mapped to more than one locus using samtools^73^; (2) were duplicated using PICARD (v2.2.4; http://broadinstitute.github.io/picard); and (3) mapped to the mitochondrial genome.

### Genotype calling

Genotypes were called by GATK (v3.5.0)^74^. We performed: (1) indel-realignment; (2) base score recalibration; and (3) joint genotype calling across all samples for variants having a phred-scaled confidence threshold ≥ 10. We excluded clustered variants, variants in ENCODE blacklisted regions^75^, and variants not present in dbSNP v146^76^. Genotype concordance between samples was assessed using both the kinship coefficient calculated by KING v1.9^77^ and the fraction of concordant genotype calls. For these analyses, we kept only variants with minor allele frequencies (MAF) ≥ 25%. The two approaches yielded comparable results, with both indicating a clear and unambiguous separation of samples. Using this approach, we were able to confirm that neuronal and non-neuronal libraries supposedly originating from the same subject showed markedly higher genotype concordance score compared to the comparison with unrelated samples (**Figure S1M**).

### Sex determination of samples

The sex of the samples was assessed using three metrics: (1) the heterozygosity rate of chromosome X genotype calls outside the pseudoautosomal regions. For this, we removed variants with MAF < 5%. A high heterozygosity rate can indicate contamination in male samples. (2) The read counts of OCRs adjacent to *FIRRE* and *XIST* genes that are predominantly expressed in females. (3) Read counts in OCRs on chromosome Y outside the pseudoautosomal region. Using this approach, we detected and excluded two samples that were supposed to originate from a male subject but they were genetically females. After their removal, all remaining samples matched the expected sex characteristics (**Figure S1L**).

### Quality control of ATAC-seq samples

For each sample, we calculated the following metrics: (1) total number of initial reads; (2) number of uniquely mapped reads; (3) fraction of reads that were uniquely mapped and additional metrics from the STAR aligner; (4) Picard duplication and insert metrics; (5) rate of reads mapped to the mitochondrial genome; (6) PCR bottleneck coefficient (PBC), which is an approximate measure of library complexity estimated as (non-redundant, uniquely mapped reads)/(uniquely mapped reads); (7) normalized strand cross-correlation coefficient (NSC) and relative strand cross-correlation coefficient (RSC), which are metrics that use cross-correlation of stranded read density profiles to measure enrichment independently of peak calling; (8) fraction of reads in peaks (FRiP), which is the fraction of reads that fall in detected peaks (see below for peak calling) and similarly the fraction of reads in only blacklisted peaks and the ratio between these two metrics. **Table S2** describes the main QC metrics. On average, we obtained more than 27 million uniquely mapped paired-end reads per sample. The rate of reads that mapped to the mitochondrial genome was below 2% since we generated ATAC-seq libraries using FANS separated nuclei, instead of whole cells. The bigWig tracks for each sample were manually inspected. A total of six libraries were excluded, having failed QC (including sex check) and/or visual inspection in IGV, leaving 70 libraries that were subjected to further analysis (**Table S2**).

### Peak calling and read quantification

First, we merged the BAM-files of samples of the same diagnosis and cell type and subsampled to a uniform depth of, at most, 454 million paired-end reads. We subsequently created bigWig files and called peaks using these merged bam files and created a joint set of peaks requiring each peak to be called in at least one of the merged BAM-files. Peaks for OCRs were called by MACS (v2.1)^78^, using the following parameters^79^: *--keep-dup all --shift −100 --extsize 200 -- nomodel.* After removing peaks overlapping the blacklisted genomic regions, 371,820 peaks remained. Next, we counted how many reads for each sample overlapped consensus peaks using the featureCounts function in RSubread^80^ (v.1.15.0). We counted fragments (defined from paired-end reads), instead of individual reads. This resulted in a sample by peak matrix of read counts, obtained using the following parameters: *allowMultiOverlap = F, isPairedEnd = T, strandSpecific = 0, requireBothEndsMapped = F, minFragLength = 0, maxFragLength = 2000, checkFragLength = T, countMultiMappingReads = F, countChimericFragments = F*.

### Differential analysis of chromatin accessibility

We performed a statistical analysis of chromatin accessibility to detect genomic regions with significant differences in chromatin structure among neuronal and non-neuronal cells. First, we used the sample-by-peak read count matrix (70 samples by 371,820 OCRs). We subsequently excluded 1,178 OCRs using a criteria of “CPM ≥ 1 in at least 10% of the samples”, resulting in our final sample-by-peak read count matrix (70 samples by 370,642 OCRs). From here, we applied the trimmed mean of M-values (TMM)^81^ to normalize the read count followed by quantile normalization to achieve a balanced distribution of reads across samples of the same cell type.

#### Covariate exploration

Next, we tested whether we could find biological or technical sample-level covariates that affect the observed read count. For these covariates (e.g. number of peaks called in the sample, FRiP, chrM metrics, RSC and NSC, and Picard insert metrics), we normalized to the median of the cell. All 63 covariates were then tested for inclusion in differential analysis as detailed in the following: As a starting point for building the model to explain chromatin accessibility in the peaks, we selected cell type by diagnosis (2×2=4 levels) and sex (2 levels). To select additional covariates, we sought a good “average model” of chromatin accessibility over all OCRs. For each additional tested covariate, we asked how many OCRs showed an improved Bayesian Information Criterion (BIC) score minus how many showed a worse BIC score when the covariate was added to the “base” linear regression model. Here, we required that at least 5% of the OCRs showed a change of 4 in the BIC score, corresponding to “positive” evidence against the null hypothesis^82^. However, no covariate satisfied the BIC score criteria for inclusion. We were unable to find any covariate even after adjusting the threshold of minimal BIC (tested values = *{2, 4, 10}*) and/or minimal fraction of OCRs exceeding this threshold (tested values = *{2%, 5%}*). Overall, our final model included 2 variables (cell type by diagnosis [4] and sex [2] ^39^), where the number of levels for factor variables is noted here in square brackets. This model accounted for 5 DF.

#### Differential analysis

We used the *voomWithQualityWeights* function from the *limma* package^41^ to model the normalized read counts. Then, we performed differential chromatin accessibility analysis by fitting weighted least-squares linear regression models for the effect of each variable on the right-hand side on accessibility of each OCR:

chromatin accessibility ∼ cell type:diagnosis + Sex + (1|Person_ID)

#### Validation of differential OCRs

To validate the relevance of differential OCRs, we applied the following strategies: permutation test and machine-learning test. For the former one, we randomly permuted MDD case/control status (*n* = 100 permuted datasets) and performed differential analysis using the same setting as for primary analysis. We measured (i) whether the sets of differential OCRs on permuted datasets are smaller compared to non-permuted datasets and (ii) whether the *P*-value rankings of differential OCRs on non-permuted datasets are close to normal distribution. For machine learning validation, we trained six machine learning models for prediction of MDD case/control status built on the reported set of (i) differential OCRs and (ii) the same number of randomly selected OCRs. We applied the repeated 5-fold cross-validation (k_repeat_ = 10) and, additionally, we repeated the whole process 10 times with different sets of randomly selected OCRs. Then, we measured an improvement of prediction performance of the classifier based on differential OCRs over classifiers utilizing random OCRs. The following machine learning methods were tested, using the default setting in R-package^83^: Naive Bayes (nb), Random forest (rf), Nearest neighbor (knn), Logistic regression (multinom), SVM with linear kernel (svmLinear), and SVM with polynomial kernel (svmPoly).

### Annotation of OCRs and gene set enrichment analysis

We determined the genomic context per each OCR based on its proximity to the closest gene as assigned by ChIPSeeker^84^. For this, we created a transcript database using GenomicFeatures^85^ and Ensembl genes. The genomic context was defined as promoter (+/- 3kb of any TSS), 5’-UTR, 3’-UTR, exon, intron, and distal intergenic. We used GREAT approach^86^ to assign OCRs to genes and perform enrichment analysis with combined set of Gene Ontology^27^, biological processes with the curated canonical pathways from REACTOME^87^, KEGG^88^, and PID^89^, all accessed from MSigDB 6.0^90^. We further pruned highly similar gene sets by iteratively removing those with a Jaccard index ≥ 0.5, preferentially keeping the bigger gene set. This resulted in 4,590 gene sets (biological processes and pathways).

### Overlap of OCRs with common variants in MDD

To determine whether the sets of neuronal, non-neuronal, and consensual OCRs as well as differential OCRs are enriched for common MDD GWAS variants^91^, we calculated partitioned heritability using LD-sc^25^. This analysis assesses if common genetic variants in the genomic regions of interest explain more of the heritability for a given trait than genetic variants not overlapping the genomic regions of interest, normalized by the number of variants in either category. The algorithm allows for correction of the general genetic context of the annotation using a baseline model of broad genomic annotations (like coding, intronic, and conserved regions). By using this baseline model, the algorithm focuses on enrichments above those expected from the general genetic context of the interrogated regions. We excluded the broad MHC-region (chr6:25-35MB) and, otherwise, used default parameters.

### Motif Matching

In order to identify candidates for DNA-binding proteins with recognition motifs enriched in our MDD-specific OCR set, we utilized the RSAT suite *peak-motifs,* a computational pipeline that discovers motifs in input sequences, and compares them with position-specific scoring matrix (PSSM) transcription factor databases^24, 92^. Input sequences are scanned to predict binding sites, and the background model is a Markov chain of order 2 trained on the input sequences. Using *peak-motifs*, word-based analysis was first performed on the MDD-specific OCR set (n=183 sequences) with hexanucleotides (*k = 6*) and heptanucleotides (*k = 7*). The tool combines four pattern-discovery algorithms that utilize overrepresentation and positional bias as two criteria to detect significant oligonucleotide, which are then used as seeds to build probabilistic description of motifs (PSSMs), indicating residue variability at each position of the motif. Discovered motifs were compared with the JASPAR nonredundant core database of known transcription factor binding motifs to predict associated transcription factors (using *compare-matrices*). Several metrics are computed to measure the similarity between each matrix pair (including Pearson correlation, width normalized correlation). These metrics are converted to ranks, and a mean rank is computed to enable comparison between candidate factors. The *peak-motifs* pipeline discovered a motif (**Figure 2A**) that was significantly enriched in MDD-specific OCR sequences. The distribution of this motif within OCR sequences is shown in **Figure 2A**, indicating a relatively higher number of sites near sequence centers. For the top 5 candidate transcription factors identified as matches to this motif, the bar graph in **Figure S2A** displays the consensus score from the Human Protein Atlas ^93^ for expression in human brain for each factor. The mRNA expression data is derived from deep sequencing of RNA (RNA-seq) from 37 different normal tissue types.

In order to characterize the functional role for the discovered motif from the *peak-motifs* pipeline, we utilized *GOMo* (v5.3.3), from the MEME-suite of tools^94^ (**Figure 2B**). This approach calculates associations between a user-specified DNA regulatory motif [expressed as a position weight matrix (PWM)] and Gene Ontology (GO) terms, by computing an association score between the (putative) targets of the input TF motif and each GO term in the GO map. An empirically generated p-value for the enrichment of the GO term is also computed for the association score for each GO term with respect to the motif, based on the rank sum test null model.

### Footprinting analysis

To determine the bound/unbound status of transcription factors in neuronal and non-neuronal cells as well as in MDD cases and controls, we performed footprinting analysis using TOBIAS (v. 0.12.4)^38^. Following the settings from our previous study^42^, we searched for the presence of 431 motifs representing 798 transcription factors (some motifs are shared due to their high similarity) in consensus OCRs of four merged BAM files representing both cell types & MDD diagnosis status. First, we ran the TOBIAS module ATACorrect to correct for Tn5 insertion bias in input BAM files, followed by TOBIAS ScoreBigwig to calculate footprinting scores across OCRs. Then, TOBIAS BINDetect combined footprinting scores with the information of transcription factor binding motifs to evaluate the individual binding positions of each transcription factor and determine whether a given position was bound by a given transcription factor or not for each condition, i.e. cell type and brain region. Finally, TOBIAS PlotAggregate was used to visually compare the aggregated footprints for select motifs.

### QPCR

In order to measure mRNA gene expression for gene targets of interest, FAN-sorted nuclei **(Figure S1R)** from human postmortem OFC tissues were prepared as described above, with addition of RNAse inhibitor in the sorting collection buffer, and pelleted for RNA extraction. Cultured human primary astrocytes (HPA) were washed with sterile PBS, scraped and pelleted for RNA extraction. For both nuclei samples and HPA cell samples, pellets were resuspended in RLT lysis buffer with 10% B-mercaptoethanol (B-ME), homogenized with a 22g needle and syringe, combined with equal volume 70% ethanol, and applied to Qiagen micro minelute column. RNA was washed, treated with DNAase, and eluted in 13ul of RNAse-free water, according to manufacturer’s instructions.

To measure *ZBTB7A* in bulk brain tissues, postmortem human OFC tissues were sectioned into 50 mg sections. Frozen mouse brains were sliced into 1mm coronal slices in a brain matrix, and 2mm OFC punches were removed. For both human and mouse tissues, sections were homogenized in Trizol (Thermo Fisher #15596026) with a motorized pestle, followed by chloroform extraction and precipitation with 70% ethanol. Samples were applied to a Qiagen micro minelute column, and RNA was washed, treated with DNAse, and eluted according to manufacturer’s instructions into 13ul RNAse-free water. For all RNA samples (derived from nuclei, cells, or brain tissues), 500ng of total RNA was utilized to synthesize cDNA using the Bio-Rad script cDNA synthesis kit (#1708891). From this reaction, 4 ng of cDNA was used to perform qPCR with PowerUp™ SYBR™ Green Master Mix (#A25742), according to the manufacturer’s instructions. Target gene CT values were averaged over 3 replicates, normalized to the reference gene (human brain – *HPRT1*, mouse brain – *Gapdh*), and the ΔΔCT was calculated. Graphs show experimental group fold change relative to controls, mean +/- SEM. Full list and sequences of primers used can be found in **Table S8**.

### Western Blot

In order to measure protein expression, postmortem human OFC tissues were sectioned into 50 mg sections. Frozen mouse brains were sliced into 1mm coronal slices, and 2mm OFC punches were removed. For both human and mouse tissues, sections were homogenized in 200ml RIPA cell lysis buffer, 1X protease inhibitor cocktail and 1X phospho-stop inhibitor using a 1ml dounce homogenizer. Following homogenization, lysates were briefly sonicated with a probe sonicator for five 1s pulses. Protein concentrations were measured using the DC protein assay kit (BioRad), and 20 ug of protein was loaded onto 4-12% NuPage BisTris gels (Invitrogen) for electrophoresis. Proteins were then fast-transferred to nitrocellulose membranes and blocked for 1 hr in 5% milk in PBS + 0.1% Tween 20 (PBST), followed by incubation with primary antibodies overnight at 4° C with rotation. The following antibodies were used: monoclonal rabbit anti-zbtb7a (Abcam # ab175918) (1:1000) for human blots, rabbit anti-Zbtb7a (Abcam #ab106592) (1:1000) for mouse blots, as well as rabbit anti-Gapdh (Abcam #ab9485) (1:10,000), and rabbit anti-H3.3 (Abcam #ab1791). After overnight primary antibody incubation, membranes were washed 3x in PBST (10 min) and incubated for 1 hr with horseradish peroxidase conjugated anti-rabbit (BioRad 170-6515, lot #: 64033820) secondary antibodies (1:10000; 1:50000 for anti-Gapdh antibody, BioRad) in 5% milk/PBST at RT. After three final washes with PBST, bands were detected using enhanced chemiluminescence (ECL; Millipore). Densitometry was used to quantify protein bands using ImageJ Software (NIH). Target protein measurements were normalized to Gapdh bands, and experimental group fold change was calculated relative to controls. Raw blots and indication of representative images utilized can be found in **Figure S6**.

### Animals

C57BL/6J mice were purchased from The Jackson Laboratory (Stock #024694). All procedures were done in accordance with NIH guidelines and the Institutional Animal Care and Use Committees of the Icahn School of Medicine at Mount Sinai.

### Male Chronic Social Defeat Stress Paradigm

In order to investigate the expression of Zbtb7a in the context of a mouse model of stress, the Chronic Social Defeat Stress (CSDS) in males was performed as described previously ^95^. Briefly, a cohort of 20 8-week old male C57BL/6J mice were randomly assigned to either the control or stress condition. Animals in the stress group underwent 10 consecutive days of a single 7-minute defeat session with an unfamiliar CD1 retired breeder male that had been previously screened for aggression towards C57BL/6J mice. Following the defeat session, the C57 mice spent 24 hours in the same cage as the CD1, separated by a perforated divider to allow for sensory contact. Control animals spent 24 hours in the same cage as a different male C57BL/6J for each day of the 10 day paradigm, separated by a perforated divider. The Social Interaction (SI) test was performed as described previously^46^. Briefly, in the first trial, the subject mouse was allowed to freely explore an arena with an empty mesh cage inside an interaction zone. In the second trial, a CD1 target was put in the mesh cage, and the mouse was again allowed to explore the arena. Trials are recorded and scored by Ethovision software: SI ratio score was calculated as (time spent in interaction with target)/(time spent in interaction zone without target). Control mice typically have scores ≥ 1.0, indicating increased time spent investigating the unfamiliar mouse. In the stress mice, scores < 1.0 are defined as “avoidant” and mice are described as stress susceptible, while scores > 1.0 are defined as “non avoidant” and the mice are described as stress resilient. In a typical CSDS experiment, approximately 30% of WT mice will segregate into the stress resilient group^96^.

### Viral Constructs

Viral vector constructs were generated as previously described^52^. Briefly, ZBTB7A overexpression plasmids (Origene Cat. #RC222759) were cloned into either a Lentiviral CMV-driven construct for use in cell culture experiments (shown in **Figure S2**) or a GFAP-GFP Adeno-associated virus (AAV) construct (Addgene plasmid #50473) for use in animal experiments (utilized in **Figure 3 and Figure 4**). Lentiviral vectors contained either a ZBTB7A-HA tagged overexpression construct or an empty vector expressing RFP. AAV vectors contained either an ZBTB7A OE construct or GFP. For AAV vectors utilized in **Figure 3**, a miRNA targeting endogenous Zbtb7a was generated using the BLOCK-iT™ Pol II miR RNAi Expression Vector Kit with EmGFP (Thermo # K493600), in addition to a scramble negative control (Thermo # K493600) (miR-neg) which forms a hairpin structure just as a regular pre-miRNA, but does not target any known vertebrate gene. Constructs were packaged into GFAP driven AAV expression vectors to generate AAV-GFAP-Zbtb7a-miR-GFP and AAV-GFAP-mir-neg-GFP. Purified plasmids were sent to GENEWIZ for sequence validation. Plasmids were sent to Cyagen Biosciences for packaging into Lentivirus or AAV6 serotype viruses at high titer (>10^12 units).

Negative control sequence without 5’ overhangs:

~~~
GAAATGTACTGCGCGTGGAGACGTTTTGGCCACTGACTGACGTCTCCACGCAGTACATTT
~~~

Oligos used for Zbtb7a KD:

NM_010731.3_1062_top:

~~~
TGCTGTAGAAGTCCAAGCCATTGCAGGTTTTGGCCACTGACTGACCTGCAATGTTGGACTTCTA
~~~

NM_010731.3_1062_bottom:

~~~
CCTGTAGAAGTCCAACATTGCAGGTCAGTCAGTGGCCAAAACCTGCAATGGCTTGGACTTCTAC
~~~

### Primary Human Astrocyte Cell Culture and LPS treatment

Primary Human Astrocytes isolated from human cerebral cortex and frozen at first passage were purchased from Sciencell (#1800) and cultured in Astrocyte medium (Sciencell #1801) on 50ug/ml coated Matrigel-coated plates (BD #354230). Cells were treated with lentivirus particles at MOI = ∼2 to overexpress ZBTB7A or RFP. Approximately 72 hours after lentivirus transduction, PHAs were treated with 2ug puromycin to positively select for cells expressing each construct. After 6 days of selection, cells were collected for molecular analyses (**Figure S2O-Q**). For testing ZBTB7A mRNA expression in inflammatory conditions, the cells were treated with either saline or LPS (Sigma Cat. # L2630) at 1 ug/ml for 8 hours, and collected for molecular analyses (**Figure S2R**).

### Primary Mouse Astrocytes

Primary astrocytes were cultured from frontal cortical dissections of mouse pups at P1, as previously described ^97^. Briefly, cortices were dissociated, and diluted in 10% Fetal Bovine Serum (OmegaSci, FB-11)/1% penicillin-streptomycin in DMEM (Gibco, 11995-065) and plated at a density of one brain per uncoated T75 flask. On DIV 1, plates were tapped to dislodge neurons and the media was changed to remove floating cells. Remaining astrocytes were maintained and grown to confluency and seeded at a density of approximately 3×106 cells/plate for subsequent experiments. Once confluent, the cells were treated with either saline or LPS (Sigma Cat. # L2630), and collected for molecular analyses (**Figure S2S)**.

### TRAP-sequencing Data

Polyribosome immunoprecipitation was performed as described ^98^. Briefly, mice were put through the CSDS paradigm, as described above, and sacrificed by rapid decapitation. Brain regions were dissected. Brain tissue was homogenized and homogenates were centrifuged to remove cell debris, and NP-40 (EMD Biosciences) and DHPC (Avanti Polar Lipids) were added, followed by another centrifugation step. The supernatant, which contains the ribosomes, was subjected to immunoprecipitation using anti-EGFP antibodies conjugated to Protein G magnetic Dynabeads (Invitrogen). The beads were washed, and RNA was extracted using Trizol reagent following the manufacturer’s protocol. RNA was further purified on RNeasy columns (Qiagen). RNA was amplified using the Ovation RNA-seq System V2 (NuGEN). Library preparation and amplification was performed by the Rockefeller University Genomic Facility, and libraries were sequenced on the Illumina HiSeq platform.

### Immunohistochemistry

Mice were anesthetized with intraperitoneal (i.p.) injection of ketamine/xylazine (10/1 mg/kg), and then perfused transcardially with ice cold phosphate buffered saline (PBS) followed by ice cold 4% paraformaldehyde (PFA) in PBS. Next, brains were post-fixed in 4% PFA overnight at 4° C and then transferred into 30% sucrose in PBS for two days. Brains were then cut into serial 40 µm coronal slices in a cryostat at −20C. Free floating slices containing OFC were washed 3x in tris buffered saline (TBS), incubated for 30 min in 0.2% Triton-X in TBS to permeabilize tissue, and then incubated for 1 hr at RT in blocking buffer (0.3% Triton-X, 3% donkey serum in TBS). Brain slices were then incubated overnight on an orbital rotator at 4 degrees C with primary antibodies. 24 hours later, brain slices were washed 3x in TBS and then incubated for 2 hrs at room temperature (RT) with a fluorescent-tagged AlexaFluor 680 secondary antibody. Brain sections were then washed 3x in TBS, incubated with DAPI (1:10000, lot #: RK2297251, Thermo Scientific 62248) for 5 min at RT, mounted on Superfrost Plus slides (Fischer Scientific) and then coverslipped with Prolong Gold (Invitrogen). Immunofluorescence was visualized using a confocal microscope (Zeiss LSM 780). For quantification of Zbtb7a overlap with Gfap, images were split into respective color channels, and we calculated the Mander’s Correlation Coefficient^99^ using the *coloc2* package (version 2.0.2) on FIJI, which performs pixel intensity correlation and statistical testing.

#### Antibodies used

##### Primary

Chicken-anti-GFAP (astrocyte marker): Thermo Scientific # PA1-10004 (1:1000)

Mouse-anti-NeuN (neuronal marker): Millipore # MAB377 (1:1000)

Note viruses utilized express eGFP (Zbt-KD/GFP and ZBT-OE/GFP) and mCherry (Gi DREADD)

##### Secondary

Goat-anti-chicken Alexaflour 680: Thermo Scientific # A32934 (1:500)

Donkey-Anti-mouse Alexaflour 680: Thermo Scientific # A32788(1:500)

Donkey-anti-Rabbit Alexaflour 568: Abcam #ab175470 (1:500)

Goat-Anti-Armenian hamster: Jackson Immunoresearch #127-545-099 (1:500)

### Animal Surgeries

Male C57BL/6J mice were anesthetized with a ketamine/xylazine solution (10/1 mg/kg) i.p., positioned in a stereotaxic frame (Kopf instruments) and 1 µl of viral construct was infused bilaterally into the OFC using the following coordinates; AP, 2.6 mm; ML, ±1.2 mm; V, 2.8 mm, angle 10°). Following surgery, mice received meloxicam (1 mg/kg) s.c. and topical antibiotic treatments for 3 days. All behavioral testing or electrophysiological recordings commenced 21 days after surgery to allow for maximal expression of the viral constructs.

### Magnetic Cell sorting

For magnetic-activated cell sorting, we collected virally-infected fresh OFC tissues, pooling 3 mice per *n*, and performed the MACs protocol, following manufacturer’s instructions. Briefly, OFC tissues were removed and washed in cold D-PBS, and tissue was dissociated using the Adult Brain Dissociation Kit, mouse and rat (Miltenyi # 130-107-677) enzyme kit in combination with the gentleMACs Octo Dissociator with Heaters (Miltenyi # 130-096-427). Samples were strained with MACs SmartStrainers (70uM, Miltenyi# 130-098-462), and spun at 300g for 10 minutes at 4C. Myelin debris was removed using myelin removal beads (Miltenyi # 130-096-733) in combination with the autoMACs Pro Separator. Samples were magnetically labeled with Anti-ACSA-2 Microbeads (Miltenyi #130-097-678) to isolate astrocytes with the autoMACs Pro Separator with the positive selection program. The negative fraction was subsequently incubated with Adult Non-neuronal Cell biotin-antibody cocktail (Miltenyi #130-126-603), followed by anti-biotin microbeads (Miltenyi #130-126-603) and then processed on the autoMACs Pro Separator to isolate neuronal cells via negative selection. For validation experiments in **Figure S2H-J**, negative fractions following astrocyte isolation were further processed with anti-Cd11b microbeads (Miltenyi #130-093-634) to isolate microglia, followed by incubation with anti-Pdgfra microbeads (Miltenyi # 130-094-543) to isolate immature oligodendrocytes, using the autoMACS Pro Separator prior to isolation of neuronal fraction, as described above. Isolated astrocyte and neuronal cells were then counted, with 50K cells separated for ATAC-seq, and the remainder of cells used for RNA extraction via trizol, followed by cleanup using the Qiagen Minelute kit. For the mouse ATAC-seq, MACs-isolated cells were processed according to the OMNI-ATAC protocol^100^, which has been optimized for fresh cells.

### Mouse ATAC-seq Differential Accessibility Analysis

Raw sequencing reads were aligned to the mouse genome (mm10) using default settings of HISAT2^101^. Only uniquely mapped reads were retained. Alignments were filtered using SAMtoolsv1.19^102^ to remove duplicate reads. Peak calling was performed using MACSv2.1.124 with settings --nomodel --shift −100 --extsize 200. Peaks were filtered for FDR < 0.05. Differential analyses were performed using diffReps20 with a window size of 1 kb. A default p-value cutoff of 0.0001 was used. Peaks and differential sites were further annotated to nearby genes or intergenic regions using the region analysis tool from the diffReps package. DiffReps outputs can be found in **Table S6**.

### Reward Sensitivity Tasks and Operant Saccharin Behavior

Animals were single housed and given restricted access to water (4h/day for 4d) before the start of the behavioral training. During the course of the experiment, mice were given access to water for 2h each day (post-session). Reward-learning training was performed as previously described^103^, with minor modifications.

The first stage of the experiment was four days of Pavlovian cue-reward association training for reinforcement with 0.2% saccharin-solution. Modular standard mouse operant chambers enclosed in light and sound blocking cubicles were used, equipped with white house lights and ventilation fans - interior dimensions: 55.69 × 38.1 × 40.64 cm; exterior dimensions: 63.5 × 43.18 × 44.45 cm; walls: 1.9 cm) (MedAssociates, Fairfax, VT). Each chamber contained two retractable levers and one central reward magazine containing a dipper calibrated to provide ∼50ul of liquid saccharin reward per each reinforcement. Each daily session was 40-min (with operant levers retracted), in which mice learned to introduce their noses into the central reward magazine to get saccharin rewards, which were delivered every 60 s. A cue light above the magazine signaled reward delivery. Correct and incorrect saccharin retrieval was detected via infrared beam breaks upon head entry in the magazine and automatically recorded by MedPC software.

Next, mice were put through 5-7 days of 1 h sessions of operant learning training, in which mice were conditioned to lever press on a fixed-ratio 1 (FR1) schedule for *ad libitum* saccharin reinforcement. The basic settings were: session onset was indicated by illumination of the house light, and extension of both active and inactive levers; one active lever response (FR1) initiated magazine-cue light illumination and subsequent reward delivery, and following retrieval a 2.5 s inter-trial interval (ITI) was initiated; the session terminated after 1 hr.

In OE studies, after lever-press training as described above, mice were further trained on a reversal learning paradigm, using two levers positioned left and right of the central liquid reward magazine. For the baseline phase, mice went through 1 session/day for 8 days of training: On FR1, a response at the correct lever initiated magazine light and reward delivery,; the session terminated after 30 min. At the reversal phase, the previously incorrect lever was now correct and *vice versa*, so that non-reward-shift behavior was required; reversal testing lasted for an additional 8 days.

### Subthreshold Social Defeat Paradigm

In order to investigate the role of Zbtb7a in stress vulnerability, we performed the Subthreshold variant of the Social Defeat Paradigm (SSDS) on a cohort of 8 week old C57BL/6J male mice that were injected with either the rAAV6-GFAP-Zbtb7a OE construct or the rAAV6-GFAP-GFP empty control vector into the OFC 3 weeks previously. Half of each virus group was randomly assigned to the stress group or control group. The stress group underwent the SSDS paradigm as described previously^104^. Briefly, the stress mice were subjected to three 5-min defeat sessions with an aggressive CD1 male mouse consecutively on a single day, separated by a 15-minute rest period. The experimental mouse then spent 24 hours in the aggressor home cage, separated by a perforated divider to allow sensory exposure to the aggressor, and was then tested for social interaction as described above. Note that WT mice do not show behavioral deficits after the SSDS paradigm.

### Forced Swim

The forced swim test (FST) was similarly conducted as previously described^52^. Mice were placed in a 4 liter glass beaker with 2L of room-temperature water for 7 minutes. Each session was recorded and scored by a blinded observer to record the number of seconds each mouse was immobile during the last 4 minutes of the test.

### Singe-cell suspension preparation and Flow Cytometry

Single-cell suspensions from the brain tissue were prepared as described previously^105^. Briefly, virally-transduced OFC tissue was dissected, minced and digested with 450 U/ml collagenase I, 125 U/ml collagenase XI, 60 U/ml DNase I and 60 U/ml hyaluronidase (Sigma) in PBS for 40 min at 37 °C. Samples were passed through a 70-μm cell strainer and mixed with 30% percoll layered on top of 70% percoll. The percoll gradient was centrifuged at 500 *g* for 30 min with the brake off. The cell fraction was collected and washed with PBS before downstream applications. Total viable cell numbers were quantified using counting beads (Thermo Fisher Scientific). Cell suspensions were stained with the antibody cocktail in PBS supplemented with 2% FBS and 0.5% BSA. The following monoclonal antibodies were used for flow cytometry analyses at a dilution of 1/700: anti-CD45 (BioLegend, clone 30-F11, 103147), anti-CD11b (BioLegend, clone M1/70, 101226), anti-CD11c (Biolegend, clone N418, 117333), anti-TREM2 (R&D Systems, clone 237920, FAB17291P), anti-P2RY12 (Biolegend, clone S16007D, 848003), anti-ASCA2 (Miltenyi Biotec, clone REA969, 130-116-245), anti-MHCII (BioLegend, clone M5/114.152, 107602) and anti-CCR2 (R&D systems, clone 475301, MAB55381). Viable cells were identified through negative staining for Zombie NIR (BioLegend). Data were acquired on a Cytek Aurora and analyzed with FlowJo (Tree Star). Flow cytometry gating strategy shown in **Figure S6** included all cells, singlets, live cells and cell populations were identified as astrocytes (ACSA2^+^CD45^−^) or microglia (CD45^mid^P2RY12^+^CD11b^+^).

### Electrophysiology

Male C57BL/6J mice (age approximately 60 days) were deeply anesthetized with isoflurane and then decapitated, followed by rapid removal and chilling of the brain. Coronal slices (300 µm thick) were prepared using a Compresstome vibrating microtome (Precisionary, Natick, MA), in ice-cold sucrose cutting solution (in mM: 215 sucrose, 2.5 KCl, 1.6 Na_2_HPO_4_, 26 NaHCO_3,_ 4 MgSO_4_, 1 CaCl_2_ and 20 glucose). The slices then were transferred to artificial cerebrospinal fluid (ACSF; in mM: 120 NaCl, 3.3 KCl, 1.2 NaHPO_4_, 26 NaHCO_3,_ 1 MgSO_4_, 2 CaCl_2_ and 11 glucose; pH 7.2, 300 mOsM; bubbled with 95% O_2_/5% CO_2_) at 32°C for 30 minutes, after which they were transferred to room temperature ACSF and allowed to recover for at least one hour. Recordings were obtained in a submersion recording chamber superfused with ACSF (1 mL/min) at room temperature. A concentric bipolar stimulating electrode was placed in layer 1 of the orbital frontal cortex to evoke synaptic responses using a 100 µs stimulus delivered by an IsoFlex stimulus isolator (AMPI, Jerusalem, Israel). A glass Ag/AgCl electrode filled with ACSF recorded field excitatory synaptic potentials (fEPSPs) from layer 5. Recordings were acquired using Axoclamp 2A and Axopatch 1D amplifiers, Digidata 1440A analog-digital convertor, and pClamp software 10 (all from Molecular Devices, San Jose, CA). Signals were low-pass filtered at 2 kHz and digitized at 10 kHz. An input-output (I-O) curve was constructed by recording fEPSPs in response to stimuli ranging from 100-800 µA (average of three fEPSPs per stimulus strength, recorded at intervals of 20 seconds between stimuli, starting with the lowest intensity). Given the proximity of the recording electrode to the stimulating electrode within respective layers of the OFC, we plotted peak amplitude (instead of peak slope), to avoid effects of recording artifacts. Following construction of an I-O curve, the stimulus intensity that evoked a fEPSP of ∼50% of maximum amplitude was used in rundown experiments. Separate slices from the same animals were used for the rundown experiments. For rundown experiments, a single 30-s train was delivered at 10 Hz after establishment of a stable baseline. The percentage change in fEPSP amplitude from baseline was calculated. All data were graphed as means ± SEM.

### Calcium Imaging

Calcium imaging was performed in 2D primary mixed cultures of mouse cortical neurons and glia (including astrocytes) to assess neuronal and astrocytic activity, using the genetically encoded calcium indicator GCaMP6f. Primary mixed cultures were transduced with AAV1-hSyn-GCaMP6f (Addgene # 100837-AAV1) or AAV5-gfaABC1D-cyto-GCaMP6fto (Addgene # 52925-AAV5) to ensure selective expression solely in neurons or in astrocytes, respectively, at least 5 days prior to the imaging sessions. Both GCaMP6f-astrocyte and GCaMP6f-neuronal cultures were treated with the AAV-GFAP-ZBTB7A OE vector for ZBT-OE conditions. In GCaMP6f-neuron cultures, to control for the astrocyte AAV treatment, “control virus” conditions were additionally treated with an AAV5-RFP empty vector. Neurons and astrocytes activity were imaged independently after 14-23 DIV, in mixed cultures plated on poly-d-lysine matrix (0.1 mg/mL, gibco #A38904-01) coated 10 mm glass coverslips. A Nikon Eclipse TE2000-U microscope with a 10X objective was used to image the coverslips with the mixed primary cultures mounted on a diamond-shaped chamber. To excite and detect GCaMP6f fluorescence, a 480 nm LED (Mic-LED-480A, Prizmatix), a HQ480/40x excitation filter, a Q505LP dichroic mirror, and a HQ535/50m emission filter (Semrock) were used. Emitted fluorescence was projected onto a sCMOS Zyla chamber camera (VSC-01910, Andor) and sampled at 8.87 fps for GCaMP6f-expressing neurons (284×240 pixels, 3×3 binning) and 4.7 fps for GCaMP6f-expressing astrocytes (160×135 pixels, 4×4 binning). Nikon Elements software (NIS-Elements AR 5.20.01) was used to control light source and sCMOS camera.

To record spontaneous neuronal or astrocytic calcium spikes, mixed cultures were continuously perfused during fluorescence recording with artificial cerebrospinal fluid buffer (ACSF), with the following composition (in mM): NaCl 125, KCl 5, D-Glucose 10, HEPES-Na 10, CaCl_2_ 3.1, MgCl_2_ 1.3. (pH adjusted to 7.4 with HCl and osmolarity corrected with sucrose to 290-300 mOsm). Perfusion was gravity fed (flow rate of 0.065 ml/s) and controlled with a ValveBank8 II (AutoMate Scientific Inc.).

GCaMP6f-expressing neurons or astrocytes ROIs were segmented, and raw fluorescence data were background corrected and extracted using Nikon Elements software. ΔF/F was calculated as (F_t_ – F_min_)/F_min_, being F_t_ = raw fluorescence at time t, and F_min_ = minimum fluorescence for the entire trace. A low-pass Butterworth filter was used to denoise the ΔF/F trace, and an adaptive iteratively reweighted Penalized Least Squares (AirPLS) based algorithm^106^ was applied to baseline correct the ΔF/F trace for drift, using R-Studio (R version 4.0.3). Spike detection was performed using a custom script in R that applied specific criteria for neuronal and astrocytic calcium events. For neurons, action potential-derived Ca^2+^ spikes had the following criteria (framerate acquisition of 8.87 fps): duration < 45 frames, rise phase >= 3 frames, fall phase >= 3 frames, rise phase <= fall phase, peak height > 4*SD (for ROIs with SD < 20), and peak height > 3*max background signal. For astrocytes, Ca^2+^ events detected fell in the following criteria (framerate acquisition of 4.7 fps): duration < 100 frames, rise phase >= 10 frames, fall phase >= 10 frames, peak height > 4*SD (for ROIs with SD < 20), and peak height > 4*max background signal. Final n of cells per condition was as follows: astrocyte-GCaMP6f (*n*=623 cells control virus saline, *n*= 559 cells ZBT OE saline, *n*=747 cells control virus LPS, and *n*=517 cells ZBT OE LPS) and neuron gCaMP6f (*n*=135 cells control virus saline, *n*= 1277 cells ZBT OE saline, *n*=238 cells control virus LPS, and *n*=1324 cells ZBT OE LPS). Statistical analysis was performed in GraphPad Prism 8.4.3.

### Chemogenetic Manipulation

In order to determine if neuronal hyperexcitability contributes to the observed behavioral effects of ZBTB7A OE, we performed the Subthreshold variant of the Social Defeat Paradigm (SSDS) on a cohort of 8 week old C57BL/6J male mice that were injected with the pAAV-hSyn-hM4D(Gi)-mCherry to express the inhibitory Gi DREADD (Addgene #50475-AAV2), in combination with either the rAAV6-GFAP-Zbtb7a OE construct or the rAAV6-GFAP-GFP empty control vector into the OFC 3 weeks previously. Half of each virus group was randomly assigned to the ZBT OE or GFP viral group. Both viral groups underwent the SSDS paradigm as described previously^107^. The experimental mouse then spent 24 hours in the aggressor home cage, separated by a perforated divider to allow sensory exposure to the aggressor. The mice were then single housed for 24 hours, and then half of each viral group was injected with either the DREADD agonist Deschloroclozapine^52^ (Tocris # 7193) at 1ug/kg in 1% DMSO or vehicle. Fifteen to twenty minutes post-injection, the mice were tested for social interaction as described above.

## Supporting information

Table S1_demo

Table S3_dac

Table S4_gsea

Table S5_deseq2

Table S6_diffreps

Table S7_stats

Table S8_primers

## Acknowledgements

We would like to thank members of the Maze and Roussos laboratories for critical readings of the manuscript. We would like to acknowledge the contribution of the Electrophysiology Core at Mount Sinai to electrophysiology experiments. This work was supported by grants from the National Institutes of Health: P50 MH096890 (I.M. and P.R.), R01 MH116900 (I.M.), R01MH109677 (P.R.), U01MH116442 (P.R.) and R01MH110921 (P.R.), F31 MH116588 and F99 NS125774 (S.L.F.).

## Data availability

Raw (FASTQ files) and processed ATAC-seq data (OCRs, and raw / normalized count matrices) have been deposited in Gene Expression Omnibus and are accessible through GEO Series accession number GSE149871. Browsable UCSC genome browser tracks of processed data are available at: https://labs.icahn.mssm.edu/roussos-lab/mdd_atacseq. Raw and processed RNA-seq data and mouse astrocyte-specific ATAC-seq data is accessible through GEO Series accession number GSE214922.

External validation sets: RNA-seq of MDD case/control postmortem human brains (GSE102556), TRAP-seq of astrocyte specific CSDS (GSE139684).

## SUPPLEMENTARY TABLES

**Table S1_demo:** Demographics of the postmortem brain cohort.

**Table S2_qc:** Quality control metrics of ATAC-seq dataset of postmortem brain cohort.

**Table S3_dac:** Differential analysis between neuronal and non-neuronal samples as well as among MDD cases and controls in ATAC-seq dataset of postmortem brain cohort.

**Table S4_gsea:** Gene set enrichment analysis for cell type and disease-specific sets of OCRS.

**Table S5_deseq2:** DESEQ2 outputs for all RNA-seq experiments.

**Table S6_diffreps:** Diffreps outputs for mouse ATAC-seq experiments.

**Table S7_stats**: Full statistical information

**Table S8_primers:** Full list and sequences of human primers used in qPCR experiments

## Supplemental Figures and Figure Legends

**Supplemental Figure 1.**
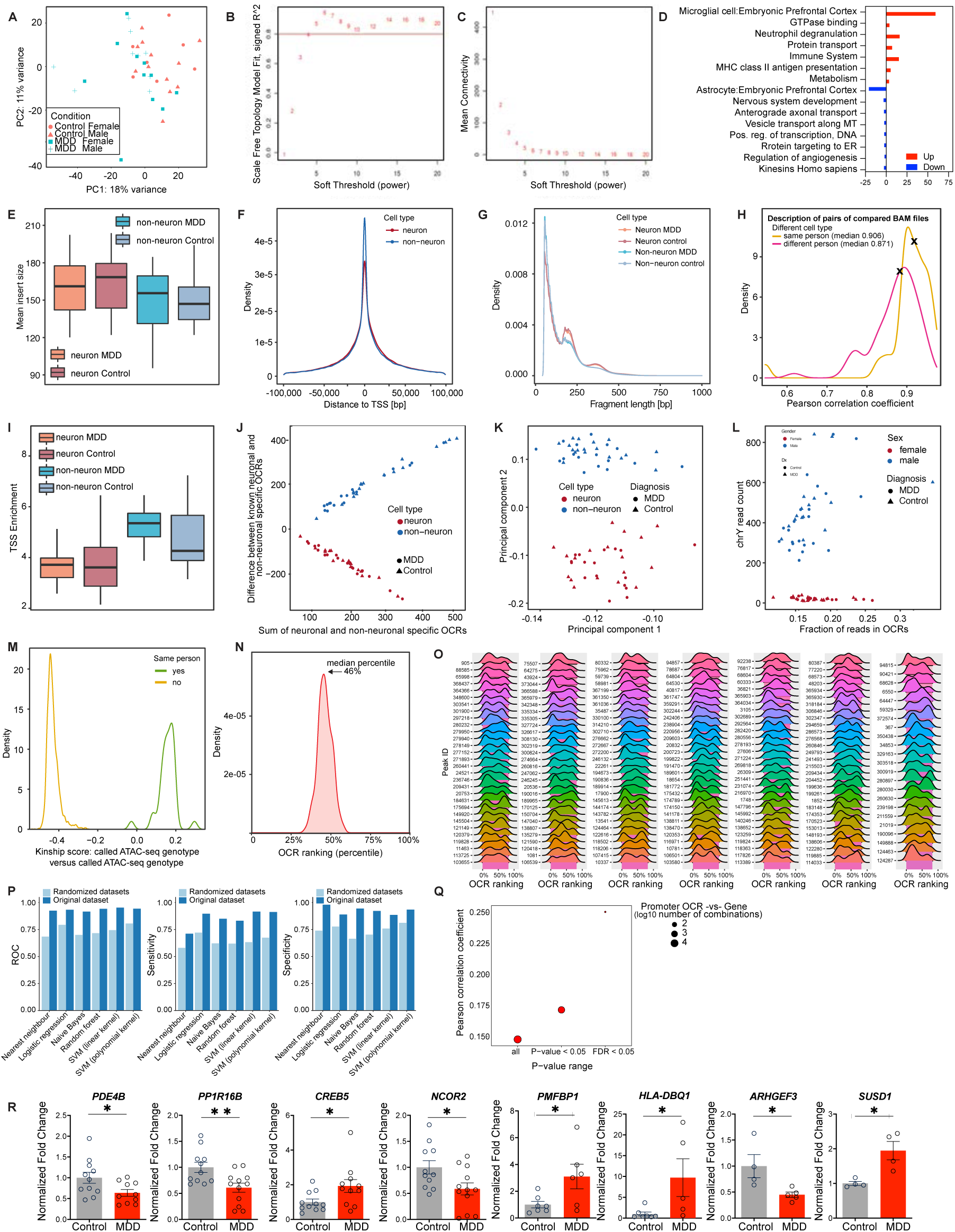
Quality control metrics for human postmortem MDD molecular profiling. (**A**) Principal component analysis of sample gene expression levels. (**B**) Analysis of scale-free fit index for possible soft-thresholding powers (β). (**C**) Analysis of mean connectivity for possible soft-thresholding powers. (**D**) GO analysis for 1,450 DE genes between MDD and control groups, separated by up/down regulation. (**E**) Fraction of uniquely mapped, non-duplicated, non-chrM paired-end reads compared to all reads in raw sequencing files. (**F**) Number of uniquely mapped, non-duplicated, non-chrM paired-end reads. (**G**) Fraction of duplicated to uniquely mapped paired-end reads. (**H**) Fraction of mitochondrial DNA reads to uniquely mapped, non-duplicated paired-end reads. (**I**) Number of OCRs (called per sample). (**J**) Fraction of reads in OCRs (FRiP). (**K**) GC-content in consensus set of OCRs; (**L**) Median insert size. For all whisker plots in this figure: The center line indicates the median, the box shows the interquartile range, whiskers indicate the highest/lowest values within 1.5x the interquartile range. (**M**) Genotype check based on pair-wise comparison of genotypes called from ATAC-seq samples. Pairs of neuronal and non-neuronal samples supposedly originating from the same person have distinctly higher scores (green line) than pairs of samples from different individuals (yellow line). (**N**) Summary and (**O**) per-OCR distribution of *P*-value ranking for the reported set of 203 differentially accessible OCRs within differentially analyses results generated on the datasets of non-neuronal samples with randomly permuted MDD and Control status (*n*=100 permuted datasets). This analysis proves that the reported set of 203 differentially accessible OCRs (median percentile of *P*-value is 1%) are not affected by technical artifacts since their median percentile of *P*-value in the datasets with permuted MDD and Control status is 46% (further details in Methods: Differential analysis of chromatin accessibility). (**P**) Performance of machine learning classifiers built on the reported set of 203 differential OCRs and 203 random OCRs. To enable the robust performance evaluation, the repeated 5-fold cross-validation was applied (*k_repeat_ = 10*); additionally, the whole process was repeated 10 times with different sets of 203 randomly selected OCRs. For all whisker plots in this figure: The center line indicates the median, the box shows the interquartile range, whiskers indicate the highest/lowest values within 1.5x the interquartile range. Student’s two-tailed t-tests were performed for statistical comparisons, *=p<.05, **=p<.01. Data displayed as mean (+/-SEM).

**Supplemental Figure 2.**
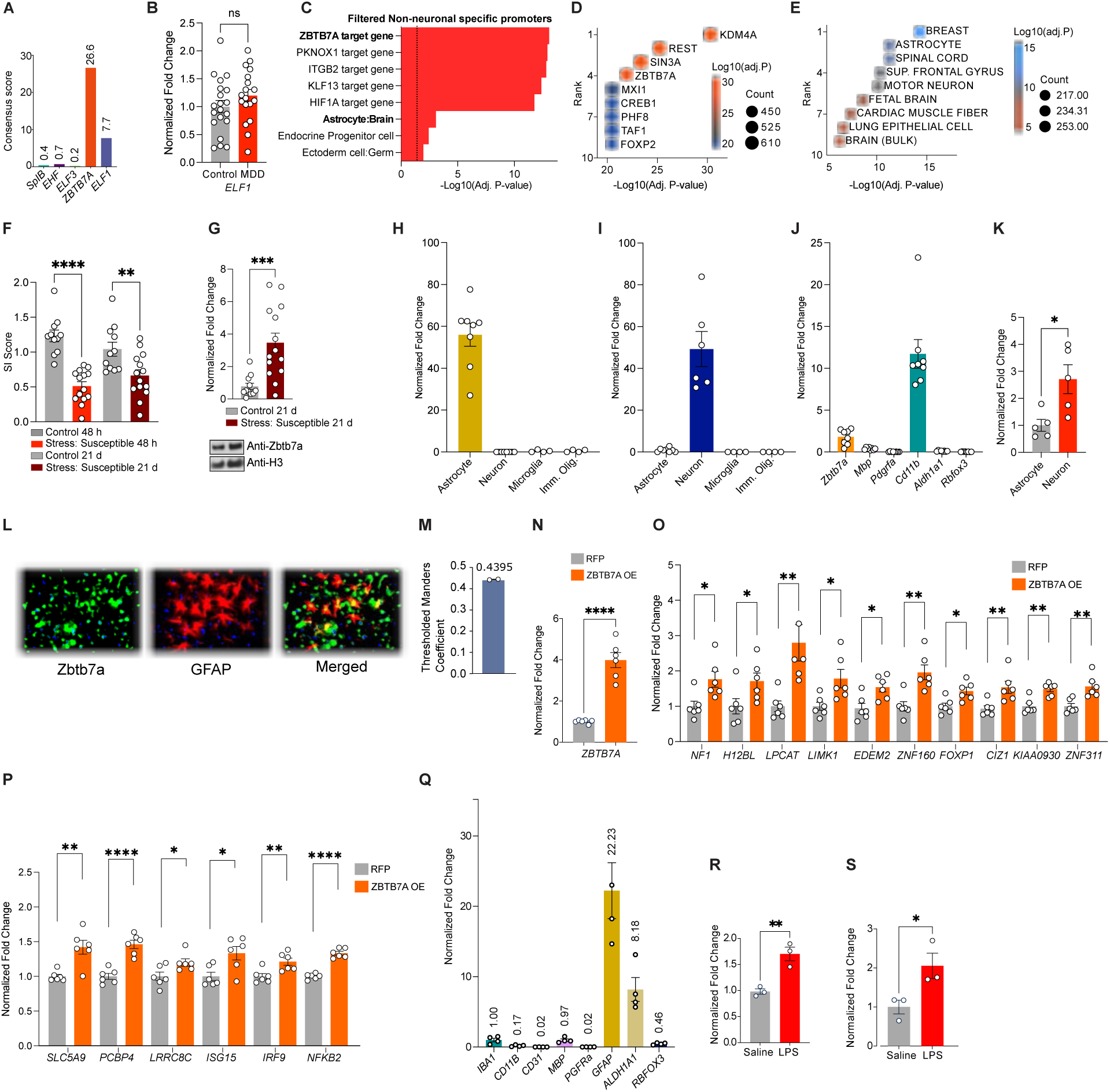
Identification and characterization of ZBTB7A in human MDD and mouse chronic stress OFC. (**A**) Consensus score from the Human Protein Atlas^1^ for expression in human brain for each factor. The mRNA expression data is derived from deep sequencing of RNA (RNA-seq) from 37 different normal tissue types. (**B**) Normalized fold change for mRNA expression for ELF1 in bulk human OFC tissues, control *vs.* MDD. (**C**) GO analysis with CellMarker Augmented Database^2^ and CHEA ENCODE Consensus database^3^ for genes in detected non-neuronal specific promoters, filtered by logFC > 1, (+/-) 3000bp from TSS (**D**) Overlap between DE genes from MDD *vs.* control OFC tissues^4^ and ENCODE consensus target gene sets via EnrichR, plotted by rank (y-axis) and - log_10_(adjusted *p*-value) on the x-axis and by fill color. Bubble size displays the number of overlapping genes for each term. (**E**) Overlap between ZBTB7A target genes (from TRANSFAC) and ARCHS4 human tissue expression reference gene sets via EnrichR. plotted by rank (y-axis) and -log_10_(Adjusted P-value) on the x-axis and by fill color. Bubble size displays the number of overlapping genes for each term. (**F**) Social interaction ratio for control vs. chronically stressed CSDS mouse groups at 48 h post-stress and 21 d post-stress. (**G**) Normalized fold change protein expression of Zbtb7a in mouse OFC bulk tissues collected from control vs. chronically stressed mouse groups at 21 d post-stress. (**H**) qPCR expression data for astrocyte-specific gene *Aldh1a1* in MACs-isolated cell fractions (**I**) qPCR expression data for neuron-specific *Rbfox3* (*NeuN* in MACs-isolated cell fractions). (**J**) qPCR expression data for cell type-specific genes in negative fraction from MACs-isolated astrocyte and neuron cell fractions, showing the negative fraction is enriched for microglia marker *Cd11b*. (**K**) qPCR expression data for *Zbtb7a* in MACs-isolated astrocyte vs. neuron cell fractions. (**L**) 20x IHC images showing Zbtb7a protein is expressed in mouse OFC astrocytes, depicts overlap of Zbtb7a with astrocyte-specific marker Gfap. (**M**) Thresholded Mander’s coefficient describes overlap of color channels of interest. (**N**) Expression of *ZBTB7A* mRNA in human primary cultured astrocytes treated with *ZBTB7A* OE lentivirus *vs.* RFP empty vector control virus. (**O-P**) Bar graph showing normalized fold change of mRNA expression in ZBT-OE vs. RFP human primary cultured astrocytes for the listed gene targets. (**Q**) Normalized fold change of cell-type specific marker genes in human primary astrocyte-enriched cultures. (**R**) Normalized fold change of *ZBTB7A* mRNA expression in cultured human astrocytes treated with saline vs. LPS. (**S**) Normalized fold change of *Zbtb7a* mRNA expression in cultured mouse astrocytes treated with saline vs. LPS. Student’s two-tailed t-tests or 1-way ANOVA with MC tests were performed for statistical comparisons. Data presented as mean (+/-SEM). *=p<.05,**=p<.01, ***=p<.001, ****=p<.0001.

**Supplemental Figure 3.**
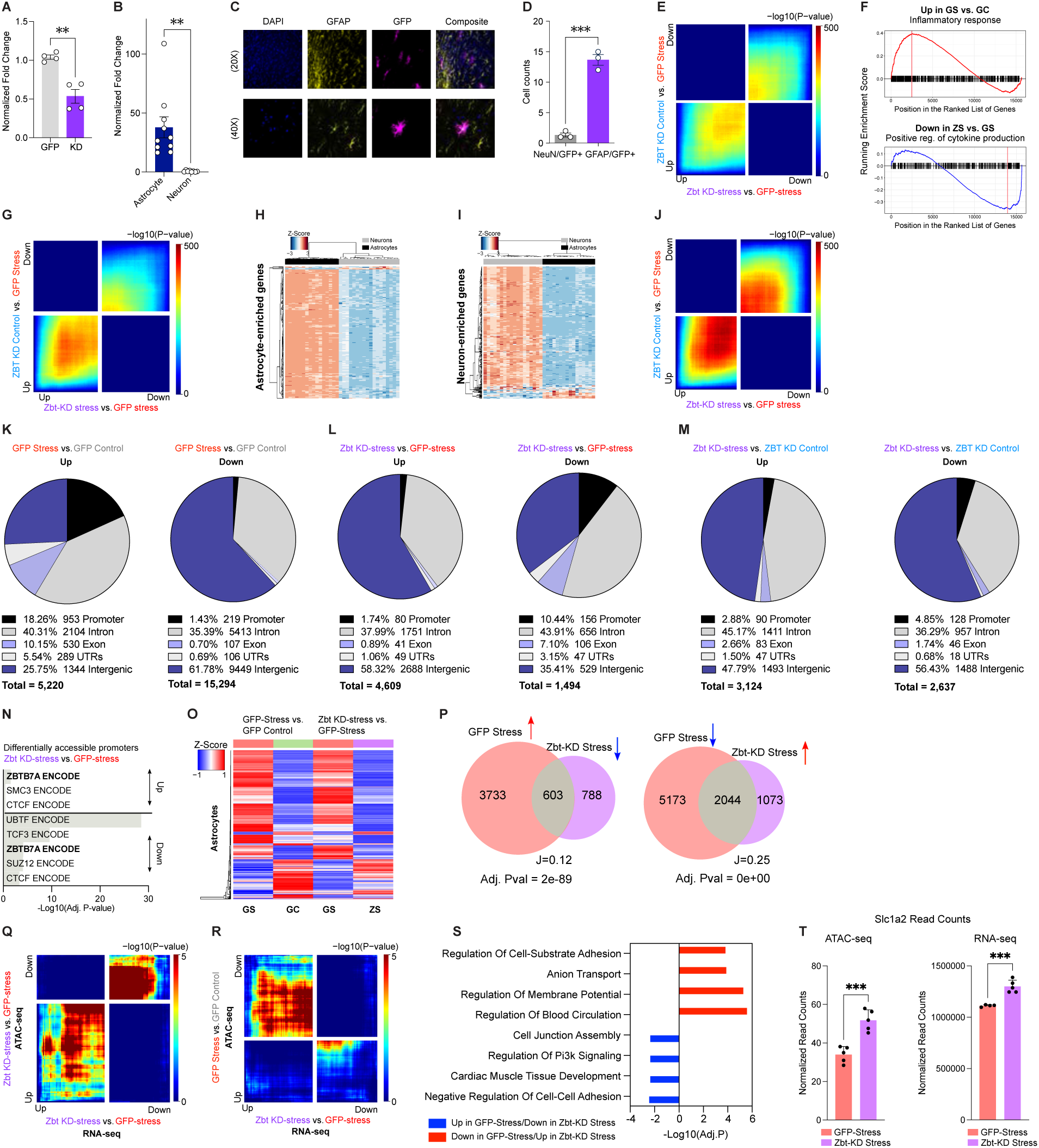
Zbtb7a KD alters cell-type specific chromatin accessibility and gene expression. (**A**) Normalized fold change of qPCR *Zbtb7a* gene expression from OFC tissues transduced with Zbt-KD virus vs. miR-neg-GFP (GFP), with n = 4/group. (**B**) qPCR expression levels of the GFP transgene in MACs-isolated neurons vs. astrocytes from AAV6-GFAP-miR-neg-GFP virally-transduced OFC mouse tissues. (**C**) Representative IHC images of OFC tissues transduced with an rAAV6 virus expressing ZBTB7A-GFP (in magenta) overlaid with a nuclear co-stain (DAPI in blue) and GFAP (in yellow) to show astrocyte-specific expression. (**D**) Cell counts in OFC tissues transduced with AAV6-ZBTB7A-GFP of cells co-expressing Gfap/Zbtb7a or NeuN/Zbtb7a. (**E**) RRHO comparing gene expression for the indicated comparisons, in bulk OFC tissue. (**F**) GSEA enrichment plot for most significantly enriched gene set in GFP Stress vs. GFP Control and ZBT stress vs. GFP Stress in bulk OFC tissue. The enrichment plot shows a line representing the running ES for a given GO as the analysis goes down the ranked list. The value at the peak is the final ES. (**G**) RRHO comparing gene expression for the indicated comparisons, in MACS-isolated astrocytes. (**H-I**) Heatmaps depict unsupervised clustering of normalized read count values in MACs-isolated astrocytes and neurons for (H) 239 astrocyte-enriched genes and (I) 279 neuron enriched genes identified in previous report^5^. (**J**) RRHO comparing gene expression for the indicated comparisons, in MACS-isolated neurons. (**K-M**) ATAC-seq diffReps analysis of differential accessibility between indicated conditions. Pie charts indicate distribution of differential accessibility events, stratified by genomic context for the indicated conditions and separated for up/down events. (**N**) Gene ontology (GO) pathway analysis of differentially accessible promoters from Zbt-KD stress vs. GFP stress [less accessible promoters, top] and GFP stress vs. GFP control [more accessible promoters, bottom]. (**O**) Clustering of groups at 1,138 overlapping genomic regions between GFP Stress vs. GFP control and Zbt-KD stress vs. GFP stress, depicting Z-score of log2FC accessibility. (**P**) Scaled Venn diagram and odds ratio analyses of the number of shared and distinct OCR gene targets between indicated conditions. Numbers indicate differentially accessible peaks, “*J*” indicates the Jaccard index. (**Q-R**) RRHO comparing gene expression and chromatin accessibility for the indicated comparisons. (**S**) GO pathway analysis of rescued OCR gene targets between Zbt-KD Stress and GFP Stress MACS-isolated astrocytes ATAC-seq. (**T**) Normalized read counts for accessibility (left) and gene expression (right) at *Slc1a2* gene in MACS-isolated astrocytes. Data were analyzed with Student’s two-tailed t-tests. *=p<.05, **=p<.01, ***=p<.001, ****=p<.0001. All data graphed as means ± SEM.

**Supplementary Figure 4.**
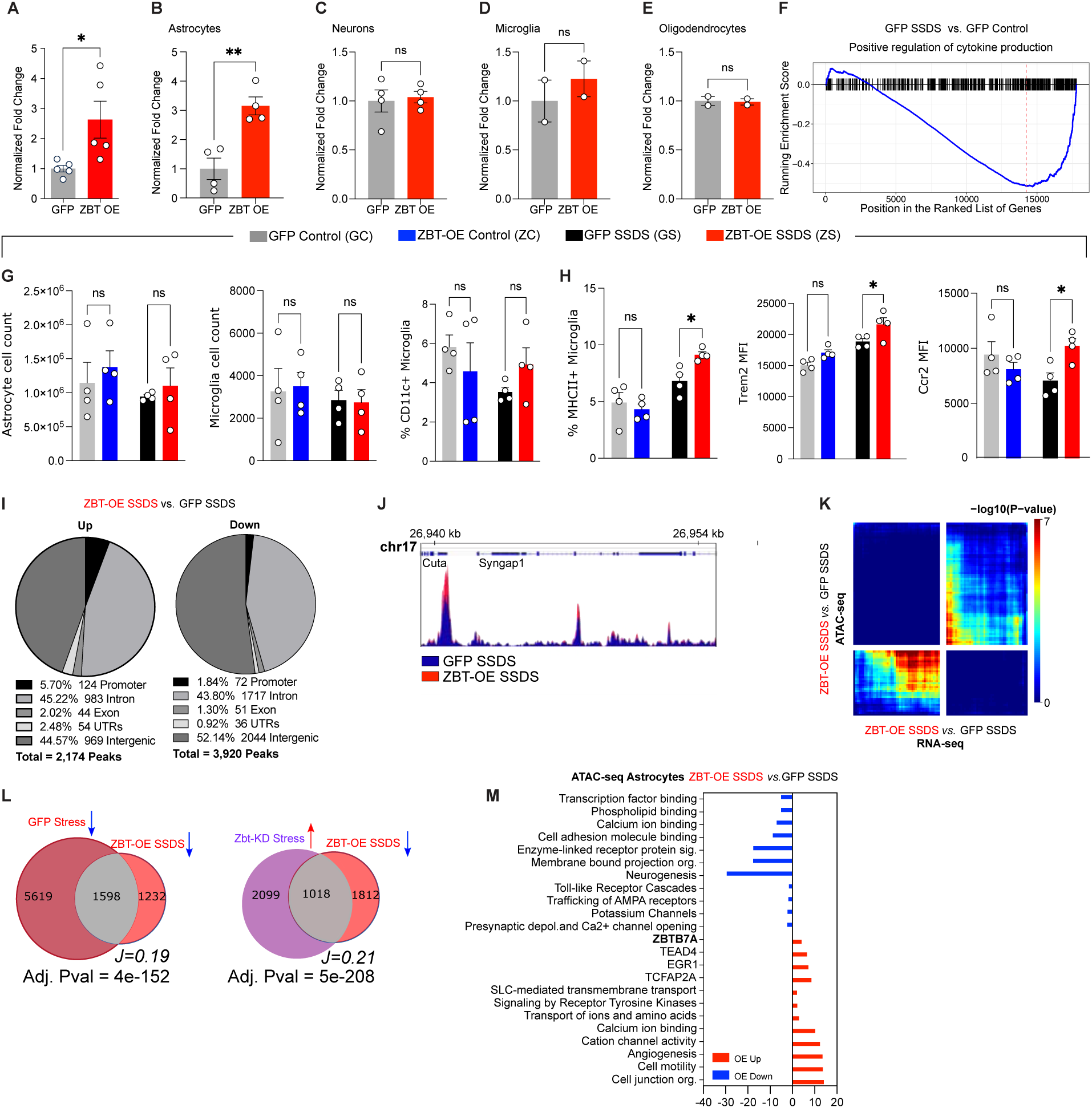
ZBTB7A OE in OFC astrocytes promotes significant alterations in behavior, chromatin accessibility, and gene expression. (**A**) Normalized fold change of qPCR *Zbtb7a* gene expression from OFC tissues transduced with ZBTB7A OE virus *vs.* GFP, n = 5/group. (**B-E**) qPCR expression levels of *Zbtb7a* in MACs-isolated (B) astrocytes and (C) neurons (D) microglia and (E) oligodendrocytes from AAV6-GFAP-ZBT OE transduced virally-transduced OFC mouse tissues, n = 2-4/group. (**F**) GSEA enrichment plot for most significantly enriched gene set in GFP Stress vs. GFP Control in bulk OFC tissue. (**G**) Number of astrocytes [left], and microglia^6^ per organ. (**H**) Percent CD11c+ microglia [far left], percent MHCII+ microglia [left], Trem2 MFI^6^ and Ccr2 [far right] MFI^6^ in virally transduced ZBT-OE vs. GFP mice (+/-SSDS) OFC via flow cytometry, n = 4/group. Gating strategy shown in **Supplementary Fig. 6.** (**I**) ATAC-seq diffReps analysis of differential accessibility comparing ZBT-OE SSDS vs. GFP SSDS. Pie charts indicate distribution of differential accessibility events, stratified by genomic context. (**J**) Representative pile-up traces of cell specific ATAC-seq signal overlapping Syngap1 gene. (**K**) RRHO comparing gene expression profile of MACs-isolated astrocytes with MACS-isolated astrocyte chromatin accessibility for indicated conditions. (**L**) Venn diagram and odds ratio analysis of the shared and distinct OCRs from ATAC-seq diffreps analysis between indicated conditions. (**M**) GO pathway analysis of gene targets associated with differentially expressed [red is more accessible, blue is less accessible] chromatin regions between ZBT-OE SSDS and GFP OE SSDS. Data were analyzed with Student’s two-tailed t-tests or with 2-way ANOVA, or 3-way ANOVA, followed by 2-Way ANOVAs for MC comparisons, *=p<.05, **=p<.01. All data graphed as means ± SEM.

**Supplemental Figure 5.**
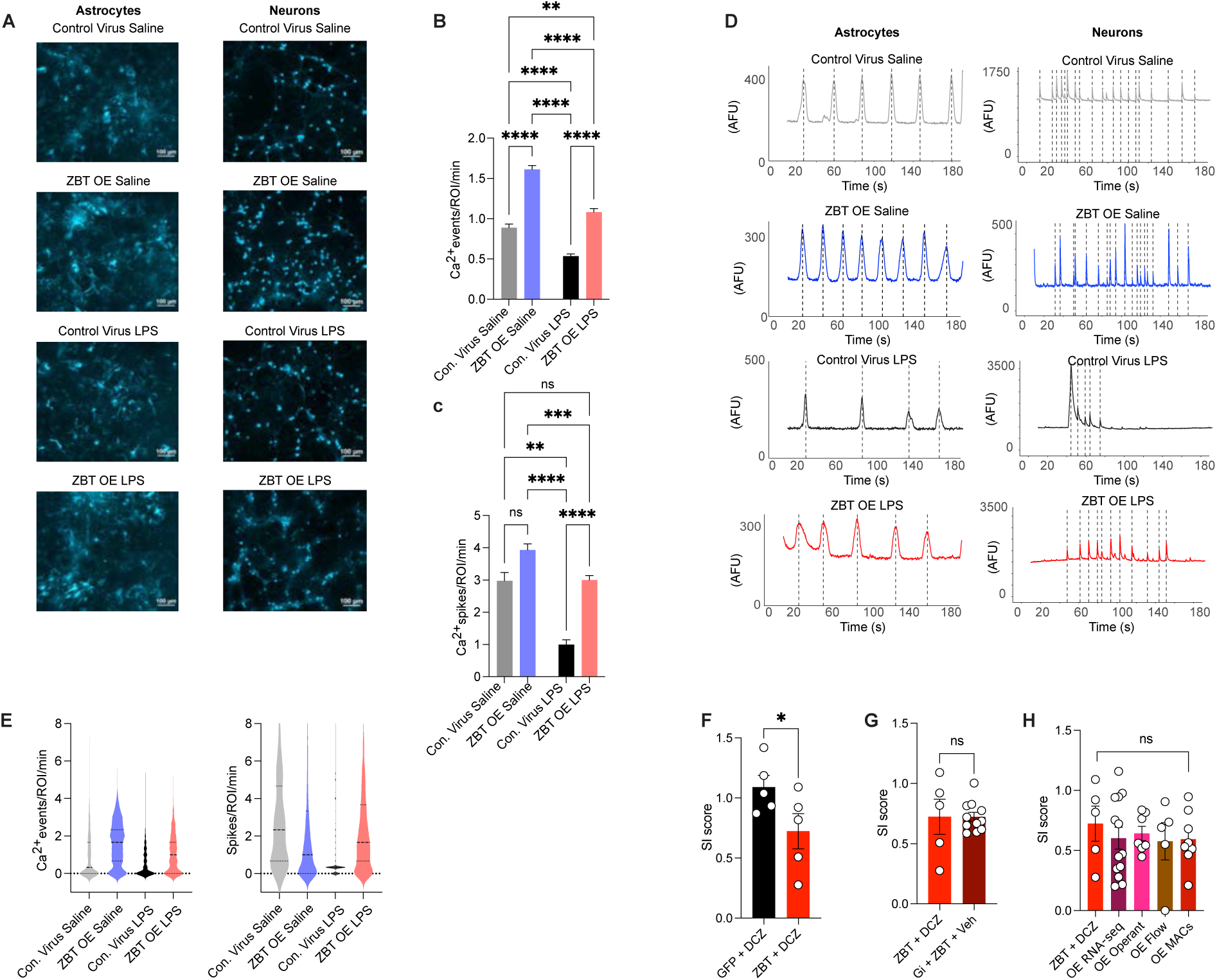
Calcium imaging and chemogenetic manipulations in the context of astrocyte-specific ZBT7A OE. (**A**) Representative images show gCAMP6f-expressing cells in either astrocyte-treated or neuron-treated primary co-cultures. (**B**) Mean frequency of Ca2+ events detected in astrocytes expressing gCAMPf. “Con.” Indicates Control. Representative traces show (**C**) Mean frequency of Ca2+ event detected in neurons expressing gCAMP6f. “Con.” Indicates Control. (**D**) Representative traces for calcium event frequencies in astrocytes [left] and neurons^6^. (**E**) Violin plots depicting individual values for (right) astrocyte [*n*=623 cells control virus saline, *n*= 559 cells ZBT-OE saline, *n*=747 cells control virus LPS, and *n*=517 cells ZBT-OE LPS] and (left) neuronal [*n*=135 cells control virus saline, *n*= 1277 cells ZBT-OE saline, *n*=238 cells control virus LPS, and *n*=1324 cells ZBT OE LPS] calcium events. (**F**) Social interaction score for ZBT OE SSDS vs. GFP SSDS mice injected with DCZ. (**G**) Social interaction score for ZBT-OE SSDS vs. ZBT-OE + G_i_ Dreadd + vehicle. (**H**) Comparison of SI score across multiple cohorts of ZBT-OE SSDS animals. Data were analyzed with Student’s two-tailed t-tests or with 1-way ANOVA plus Tukey’s MC test, *=p<.05, **=p<.01, ***=p<.001, ****=p<.0001. All data graphed as means ± SEM.

**Supplemental Figure 6.**
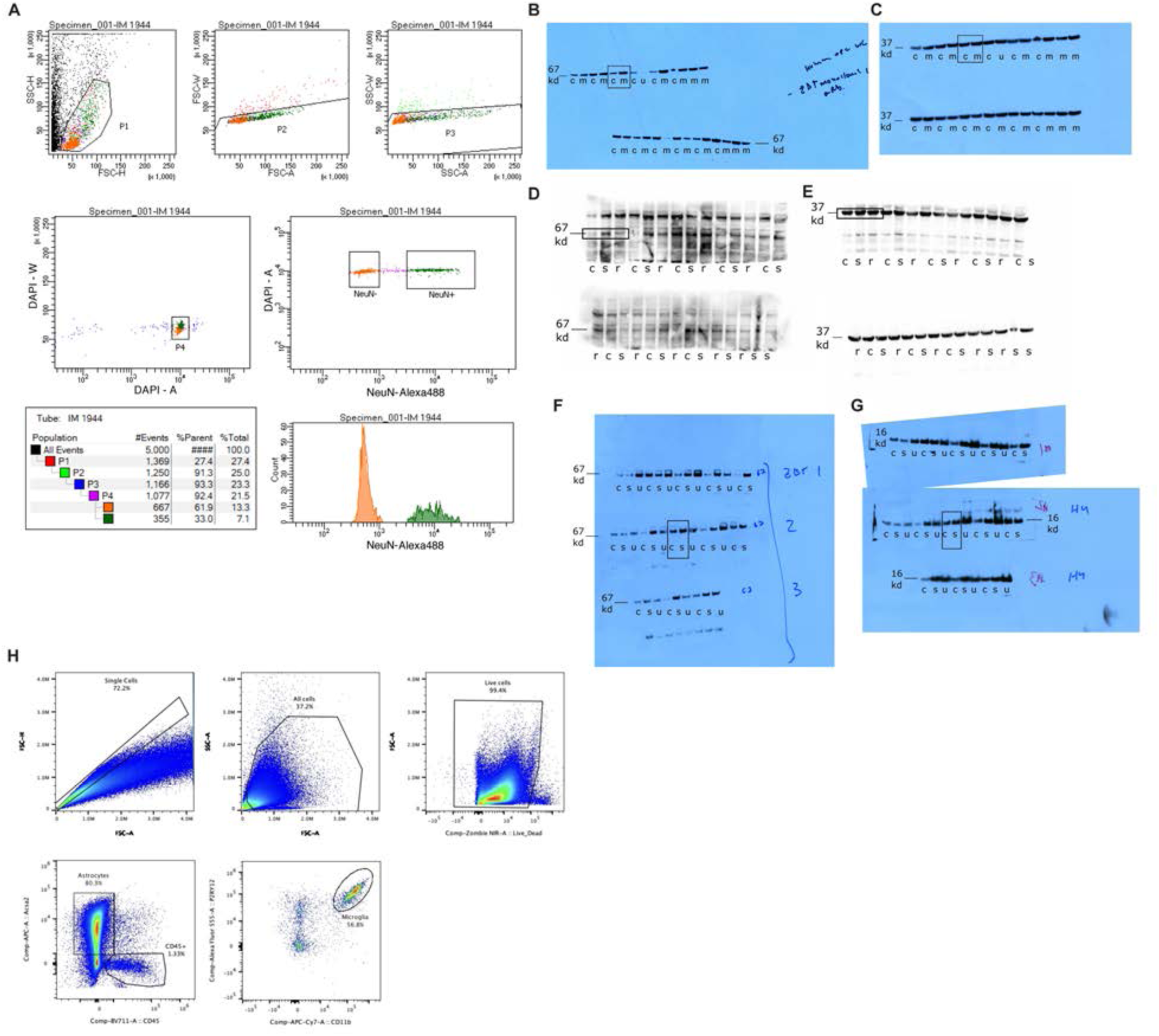
Flow cytometry gating and raw blots. (**A**) For FANS-coupled ATAC-seq on human postmortem tissues, nuclear populations were initially gated by side and forward scatter to differentiate nuclei from cellular debris. Populations were then gated based on DAPI staining to identify singlets and to further disregard debris. DAPI positive nuclei were subsequently gated based on NeuN staining to differentiate neurons (NeuN^+^) from non-neurons (NeuN^-^). Final nuclei population abundance for non-Neurons (NeuN^-^): 70.5% (in orange) and for neurons (NeuN^+^): 29.5% (in green). (**B**) Western blot film scan for ZBTB7A in bulk human OFC tissue, MDD (labeled “m”) vs. controls (labeled “c”). ZBTB7A band at expected molecular weight of 67kDa. Note samples labeled “u” are not included in this manuscript due to lack of signal (suspected improper nuclear lysis). (**C**) Western blot film scan for housekeeping gene GAPDH in human OFC, MDD vs. controls. Run on the same membrane as ZBTB7A in (B). (**D**) Raw image from chemidoc for western blot film for Zbtb7a in male mouse OFC, 48 hours after final defeat. CSDS susceptible (labeled “s”) vs. CSDS resilient (labeled “r”) vs. controls (labeled “c”). (**E**) Raw image from chemidoc western blot film for Gapdh loading control in male mouse OFC, CSDS susceptible vs. resilient vs. controls. Run on the same membrane as Zbtb7a in (D). (**F**) Western blot film scan for Zbtb7a in male mouse OFC, 21 days after final defeat. CSDS susceptible (labeled “s”) vs. controls (labeled “c”). Note samples labeled “u” are from an unrelated study, and not included in this manuscript. (**G**) Western blot film scan for H3.3 loading control in male mouse OFC, CSDS susceptible *vs.* controls (note H3.3 was used for these blots due to use of nuclear lysates, Gapdh could not be used). Run on the same membrane as Zbtb7a in (F). (**H**) Gating strategy used to identify cell populations in the OFC of mouse OE experiments (Fig. S4).

